# Southern South American Maize Landraces: A Source of Phenotypic Diversity

**DOI:** 10.64898/2026.01.02.697242

**Authors:** Tatiana Luján Dudzien, Damián Freilij, Raquel Alicia Defacio, Mariana Fernández, Norma Beatriz Paniego, Verónica Viviana Lia, Pia Guadalupe Dominguez

## Abstract

Maize (*Zea mays* L.) is a globally important crop for food, feed, and industrial uses. Modern breeding increasingly targets traits beyond yield, including stress tolerance, nutritional quality, and pest resistance. Progress toward these goals is constrained by the narrow genetic diversity of commercial varieties, a consequence of the repeated use of a limited number of inbred lines. Maize landraces therefore represent valuable reservoirs of genetic and phenotypic diversity.

Northern Argentina is one of the southernmost regions of maize landrace cultivation and comprises two main centers of diversity: Northeastern Argentina (NEA; <2000 m.a.s.l.) and Northwestern Argentina (NWA; >2000 m.a.s.l.). Despite their potential relevance, phenotypic characterization of these landraces remains limited, particularly for biochemical traits, which, although less visible, play key roles in biomass accumulation, defense against pathogens and herbivores, tolerance to environmental stress, and quality attributes such as flavor.

Here, we evaluated 17 phenotypic traits, including morphological traits, leaf biochemical compounds (such as pigments, carbohydrates, and phenolics), and salt stress tolerance, in 19 maize landrace accessions from Northern Argentina. Substantial variation was detected across all traits, both within and among accessions, indicating that each accession harbors a distinct phenotypic profile. While no significant differences were observed between regions, redundancy analysis revealed associations between phenotypic variation and collection-site altitude.

These findings highlight the value of Argentine maize landraces as sources of biochemical and stress-related traits and support their conservation and use in breeding programs aimed at broadening the genetic base of cultivated maize.

**Main Conclusion:** Maize landraces from Southern South America show high within and between accession phenotypic variability, while differences between regions of origin are associated with altitude.

## Introduction

Maize (*Zea mays* L.) is one of the most important crops worldwide, serving as a staple food for humans and livestock and as a raw material for numerous industrial products. Despite a global grain production of approximately 1.23 billion metric tons in 2024/2025 (USDA-FAS, 2025), the growing human population and increasing demand for animal-derived products continue to drive the need for higher maize yields (Andorf et al. 2019). Moreover, modern breeding goals extend beyond yield improvement to include enhanced tolerance to abiotic stresses, improved nutritional quality, pest resistance, and suitability for industrial uses (Liu et al. 2020; Prasanna et al. 2020). One of the main limitations in maize breeding is the narrow genetic base of commercial varieties, a consequence of the recurrent use of a limited set of ancestral races in breeding programs (Hufford et al. 2012; Smith et al. 2017). Indeed, commercial maize is estimated to contain only 5-10% of the total genetic diversity available in the species (Hoisington et al. 1999).

Maize was domesticated in Mexico approximately 9000 years before present (BP) (Matsuoka et al. 2002; Piperno et al. 2009) and introduced into South America around 8000 years BP (Piperno 2011; Bonavia 2013; Aceituno and Loaiza 2014). As it spread across the continent through human-mediated exchange, maize underwent substantial demographic and selective changes that led to the development of numerous local varieties, known as landraces. The term *landrace* refers to populations or groups of individuals sharing common morphological, ecological, and genetic characteristics linked to their cultivation history, which distinguish them as a recognizable group (Anderson and Cutler 1942). In general, landraces are characterized by a shared historical origin, high genetic diversity, local adaptation, identifiable phenotype, absence of formal breeding, and association with traditional farming systems (Camacho Villa et al. 2005; Guzzon et al. 2021).

Northern Argentina constitutes one of the southernmost regions for maize landrace cultivation. This area has been proposed as a historical zone of interaction between Andean and tropical lowland maize varieties (Vigouroux et al. 2008; Tenaillon and Charcosset 2011). It harbors around 57 distinct maize landraces grouped into two major genetic clusters corresponding to contrasting agroecosystems: the Northwestern (NWA) and Northeastern (NEA) Argentine maize groups (Bracco et al. 2012; Melchiorre et al. 2017; Realini et al. 2018; López et al. 2021; Rivas et al. 2022; Dominguez et al. 2024a). NWA is a mountainous region reaching up to 4000 m above sea level, characterized by low precipitation, high solar radiation, and large daily temperature amplitudes (Rivas et al. 2022). Entisols, alfisols, mollisols, and rock outcrops are the predominant soil types (Panigatti 2010). In contrast, NEA lies near sea level, with subtropical temperatures and high annual rainfall (Heck et al. 2020), and oxisols/ultisols, alfisols, mollisols, and vertisols as predominant soil types (Panigatti 2010).

Assessing phenotypic variation in crops is fundamental for identifying traits relevant to breeding (Geiler-Samerotte et al. 2013). Such variation encompasses not only morphological differences but also physiological, biochemical, and stress-related traits. Among these, resistance to abiotic and biotic stress is increasingly important under changing climate conditions. Biochemical traits, though less visible, play a key role in agronomically important processes, including biomass accumulation, defense against pathogens and herbivores, tolerance to environmental stress, and organoleptic properties such as flavor (Pott et al. 2019). Phenotypic variation in any population arises from both genetic and environmental factors. While morphological, biochemical and stress-related traits are environmentally influenced, several studies have demonstrated that their underlying genetic basis in maize is important (Reynolds et al. 2005; Mladenović et al. 2014; Kumar et al. 2015; Das et al. 2019; Santiago et al. 2025).

Maize landraces from Argentina have been extensively characterized at the morphological and phenological levels (Cámara-Hernández et al. 2012; Melchiorre et al. 2017, 2020; Rivas et al. 2022; Realini et al. 2023; Defacio et al. 2025). However, studies examining other phenotypic dimensions, such as biochemical traits, remain limited. For example, Heck et al. (2019) reported wide variation in oil and fatty acid content among 16 NEA landraces, while a correlation between amylose content and grain hardness parameters was identified in 12 landraces (Robutti et al. 1999). Mansilla et al. (2021) found differential carbohydrate and phenolic levels in two open-pollinated populations derived from Argentine landraces. Similarly, Sampietro et al. (2013) identified variation in phenylpropanoid levels associated with biotic stress resistance in three Argentine landraces. Together, these findings suggest substantial biochemical diversity within local maize varieties, underscoring their potential value for breeding applications.

In this context, the present study aims to expand the morphological, biochemical and salt-stress tolerance characterization of maize landraces from Northern Argentina to assess the extent of variability among them. To this end, 19 landrace accessions, collected from the NWA and NEA regions and conserved in the Germplasm Bank Collection of the National Institute of Agricultural Technology (INTA), were evaluated under common garden conditions. The analyses included morphological and agronomically relevant biochemical leaf traits -such as phenolic compounds, total sugars, and pigments-as well as assessments of seedling salt stress tolerance under three different salt conditions. The results provide new insights into the phenotypic diversity of Argentine maize landraces, contributing to their conservation and to breeding efforts aimed at broadening the genetic base of cultivated maize.

## Materials and Methods

### Plant Material

A set of 19 maize landrace accessions from Argentina was obtained from the “Banco Activo de Germoplasma INTA Pergamino” (BAP; Active Germplasm Bank of the National Institute of Agricultural Technology, Pergamino, Buenos Aires, Argentina), representing two distinct geographical regions, NWA and NEA (Figure 1; Supplementary Table 1).

**Figure 1.**
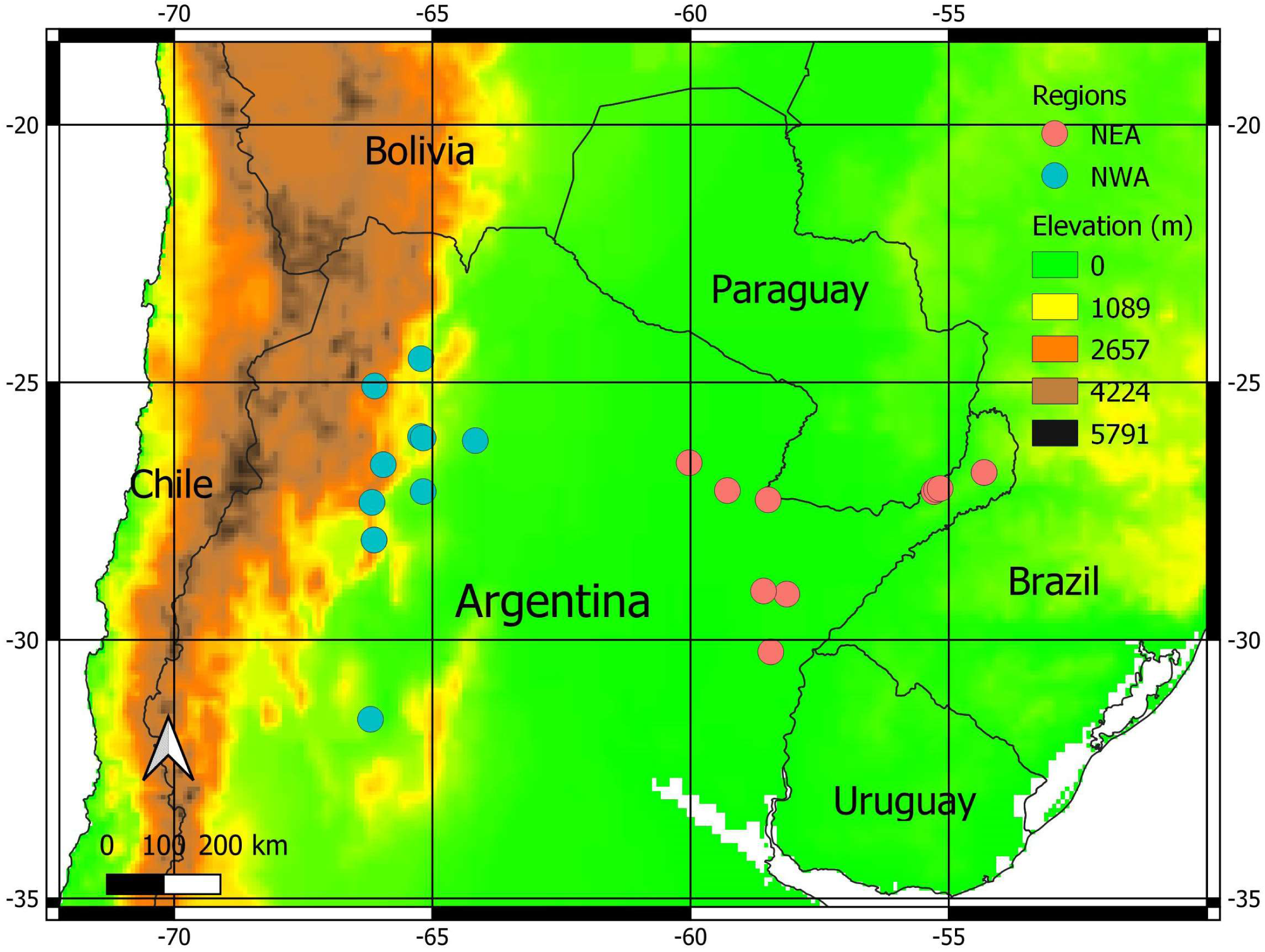
Collection sites of the accessions evaluated in this study. The list of accessions is in Table S1. NEA: Northeastern Argentina. NWA: Northwestern Argentina.

### Cultivation Conditions

Plants were grown in a greenhouse (80% relative humidity, 200 µmol PAR s⁻¹m⁻², 16 h light/8 h dark) using a completely randomized design. Five individuals per accession were cultivated in each of two independent experiments: Experiment 1 (January-April 2023) and Experiment 2 (October 2023-January 2024).

### Agromorphological Trait Analyses

Leaf number, plant height, and chlorophyll content (Dualex® optical leaf clip meter, ForceA) were measured 30- and 37-days post-sowing in Experiments 1 and 2, respectively. Because these traits were measured at different growth points, they were analyzed independently for each experiment (see statistical analysis section). Stem diameter and dry biomass (oven-dried at 80°C for 3 days) were recorded for each plant at the end of each experiment at the same growth stage (R5) and were analyzed collectively (see statistical analysis section).

### Sample Collection for Biochemical Analyses

Fully expanded leaves from V12-stage plants were collected in each experiment, immediately frozen in liquid nitrogen, stored at −80°C, and ground to a fine powder under liquid nitrogen. Data from both experiments were analyzed collectively (see statistical analysis section).

### Spectrophotometric Measurements

All biochemical measurements were performed using a Thermo Scientific Multiskan SkyHigh spectrophotometer.

### Total Protein Measurement

Protein extraction was performed by adding 1 mL extraction buffer (100 mM NaCl, 50 mM Tris pH 7.5-8, 10 mM DTT -dithiothreitol-, 20% v/v glycerol, 2 mM PMSF-phenylmethylsulfonyl fluoride) to 100 mg ground tissue. Protein concentration was determined using the Bradford assay (Bradford, 1976) adapted for microplates (He, 2011). Each well contained 200 µL of 1:4 diluted Bradford reagent (Biorad, catalogue #500-0006) and 4 µL of sample. A BSA calibration curve (0.25-10 µg/µL) was included, and absorbance was measured at 600 nm.

### Chlorophyll and Carotenoid Measurement

Pigments were extracted by adding 250 µL of 95% ethanol to 10 mg of sample and incubating at 80°C for 20 min. The supernatant was transferred to a clean tube, and the process was repeated twice to obtain pooled extracts. Absorbance was recorded at 664.2, 648.6, and 470 nm, and pigment concentrations were calculated using Lichtenthaler’s equations (Lichtenthaler 1987).

### Total Sugar Measurement

Total soluble sugars were quantified following Laurentin and Edwards (2003) using the same ethanolic extract as for pigment analysis. A glucose calibration curve (0.025-1 mg/mL) was used. The reaction was performed with 40 µL of sample and 100 µL of 0.2% anthrone reagent (Biopack, catalogue #9939.02), vortexed, incubated at 90°C for 17 min, cooled on ice, and measured at 620 nm.

### Starch Measurement

Starch content was quantified following Smith and Zeeman (2006) with modifications. The ethanol-insoluble pellet from pigment and sugar extractions was resuspended in 400 µL of 0.1 M NaOH, incubated at 95°C for 30 min, and neutralized with 80 µL of 0.1 M sodium acetate buffer. Aliquots (40 µL) were incubated overnight at 37°C with shaking in 100 µL of enzyme solution containing α-amylase (0.5 U) (Sigma-Aldrich, catalogue #A3176) and amyloglucosidase (0.45 U) (Sigma-Aldrich, catalogue #ROAMYGL) in 50 mM acetate buffer. Control samples received buffer only. The glucose released was quantified in the supernatant using the anthrone method as described for total sugars.

### Total Phenolic Compounds Measurement

Total phenolic content was determined following Ainsworth and Gillespie (2007) using the Folin-Ciocalteu assay with a gallic acid calibration curve (0.05-1 mM). For extraction, 1.5 mL of 95% methanol was added to 20 mg of sample and shaken at 4°C for 2 h. The supernatant was reacted with 200 µL of 10% Folin-Ciocalteu reagent (Biopack, catalogue #0891.05) for 6 min, followed by 800 µL of 700 mM sodium carbonate and incubation at 45°C for 15 min. Absorbance was measured at 765 nm.

### Antioxidant Activity Measurement

Antioxidant activity was determined using the ABTS assay (Re et al. 1999). For extraction, 1 mL of hexane:acetone:ethanol (2:1:1) was added to 20 mg of sample and shaken for 15 min. After addition of 150 µL of water and further mixing, the hydrophobic (upper) and hydrophilic (lower) phases were separated. ABTS (Abcam, catalogue #ab142041) was diluted in ethanol to an absorbance of 0.7, and reactions were carried out with 10 µL of extract and 200 µL of ABTS solution. A gallic acid calibration curve (0.025-0.0025 mM) was used, and absorbance was read at 734 nm in both phases.

### Salinity Stress Test

Accessions were evaluated under three NaCl treatments: 0 mM (T0), 175 mM (T1), and 350 mM (T2). Seeds (20 per accession) were sown in sand moistened with the corresponding NaCl solution and incubated for 14 days in germination chambers under the conditions outlined in the ISTA (International Seed Testing Association, 2026) guidelines: 8 hours of light at 30°C, followed by 16 hours of darkness at 20°C. Seedlings were harvested, washed, and measured for root length (from stem base to longest root tip) and shoot length (from root insertion to leaf apex). Roots and shoots from the same accession were pooled, dried at 65°C for 48 h, and weighed together to obtain a single root and shoot dry weight per accession. Three independent replicates of the experiment were performed.

### Univariate Statistical Analyses

Morphological and biochemical traits in adult plants were fitted with linear mixed-effects models to assess differences at two hierarchical levels: (1) Region, and (2) Accession. For the regional analyses (1), region was treated as a fixed factor, while accession (nested within region) and experiment were included as random effects: Y ∼ Region + (1∣Region/Accession) + (1∣Experiment). Models were fitted using the lmer() function from the lme4 package (Bates et al. 2015). For the accession-level analyses (2), accession was modeled as a fixed effect and experiment as a random effect: Y ∼ Accession + (1∣Experiment). These models were fitted with the lme() function from nlme (Pinheiro et al. 2025), allowing for heterogeneous variances among accessions or experiments using the varIdent() or varPower() variance structures when appropriate. For the three phenotypic traits measured at different plant stages in each experiment (plant height, leaf number, and chlorophyll content measured with Dualex®), analyses were conducted separately for each trial. For continuous variables (height and chlorophyll), we applied linear models to each dataset as before, with the term “Experiment” being excluded from the models. For count data (leaf number), a Conway-Maxwell-Poisson distribution (family = compois) was applied using glmmTMB (Brooks et al. 2017).

Variables associated with salt stress in seedlings were analyzed using linear mixed-effects models. Two analytical approaches were employed: (1) Region and Treatment were considered fixed factors, while Accession and Experiment Repetition were included as random factors; (2) Accession and Treatment were considered fixed factors, while Experiment Repetition was considered a random factor. Leaf and root length were analyzed using both approaches, while leaf and root dry weight were analyzed only with approach (1), as individual accession-level data were not available. In approach (1), the region × treatment interaction was initially tested but found non-significant, and models without the interaction term were subsequently fitted. The general model structure was: Y ∼ Region + Treatment + (1∣ Region/Accession) + (1|Repetition). Models were fitted using the lmer function from the lme4 package (Bates et al. 2015). In approach (2), a significant accession × treatment interaction was detected; therefore, models including the interaction term were fitted as follows: Y ∼ Accession×Treatment + (1|Repetition). Models were fitted using the lme function from the nlme package (Pinheiro et al. 2025), applying appropriate variance structures to account for heteroscedasticity among groups.

For all the analyses, model selection was based on Akaike’s Information Criterion (AIC) and inspection of diagnostic plots. Normality was evaluated with Q-Q plots and homoscedasticity with residuals versus fitted values plots. Logarithmic transformations were applied when necessary to meet model assumptions. Estimated marginal means (EMMs) were obtained with the emmeans package (Lenth and Piaskowski 2025), and pairwise comparisons were adjusted for multiple testing using Tukey’s method. Back-transformations to the original scale were performed with the “response” option. Statistical significance was set at p < 0.05.

### Multivariate Statistical Analyses

Coefficients of variation (CV) and Spearman correlations were computed for the 17 measured traits in Experiment 1 and 2 in adult plants using the *stats* and rcorr functions of the *Hmisc* package, respectively (R Core Team, 2023). Pairwise comparisons were adjusted for multiple testing using the sequential Bonferroni correction applied by the p.adjust function of *stats*. For Accession-level analyses, data were averaged per accession and then standardized and centered within each experiment to avoid potential scale-related biases. Principal Component Analysis (PCA) was conducted with the prcomp function (*stats*), and results were visualized using the fviz_pca_ind and fviz_pca_var functions from *factoextra* (Kassambara and Mundt 2020). The structure of the morphological and biochemical data was further examined through Discriminant Analysis of Principal Components (DAPC) with *adegenet* (Jombart 2008), which identifies clusters by maximizing between-group variation while minimizing within-group variation (Jombart et al. 2010). Up to 10 clusters were evaluated with find.clusters(), and cross-validation (xvalDapc, 90% training set) was used to determine the number of PCs (Principal Components) to retain, selecting the value that minimized the mean squared error (MSE). Based on this, the optimal number of groups was chosen according to the Bayesian Information Criterion (BIC). Cluster assignments were visualized with scatterplots of the first discriminant function and membership probability barplots (compoplot). Additionally, hierarchical cluster analysis was performed on adjusted means derived from the linear models performed previously. Euclidean distances between accessions were computed, and the agglomerative coefficient, calculated with agnes() (*cluster*, Maechler et al. 2022) across different linkage methods (average, single, complete, Ward’s), indicated Ward’s minimum variance method as the most suitable clustering approach. The optimal number of clusters was determined with the average silhouette method using the fviz_nbclust function (*factoextra*), and the resulting dendrogram was generated with the *ggdendro* package (de Vries and Ripley 2024). Finally, to explore potential geographical patterns, we performed a Redundancy Analysis (RDA) to quantify the proportion of multivariate trait variation (employing adjusted means derived from the univariate linear models) explained by Altitude, Latitude and Longitude (Supplementary table 1) as environmental predictors. The analysis was conducted using the *vegan* package (Oksanen et al. 2022), fitting one model with Altitude as the sole predictor and another also including Latitude and Longitude. Significance was assessed through permutation tests with 999 replicates.

## Results

### Accessions Show High Variability with Low Coefficients of Variation at the Morphological Level

Plants were evaluated in two independent experiments at the morphological level (Tables 1 and 2, Supplementary table 2). All traits showed higher CV (coefficient of variation) in Experiment 1 in comparison to Experiment 2, indicating a higher variability in the former assay (Supplementary figure 1) probably due to environmental differences between experiments combined with the high plasticity of maize.

**Table 1.** Height, Leaf number, and Total Chlorophyll measured with Dualex® in Experiment 1 (30 Days After Sowing) and Experiment 2 (37 Days After Sowing). Data represent Estimated Marginal Means or EMM (Standard Errors or SE) from Linear Mixed Effects Models, with different letters indicating significant differences (p<0.05), and Coefficients of Variation (CV). N=3-5.

| Accession | Height (cm) | | | | Leaf Number | | | | Total Chlorophyll Dualex® ($\mu\text{g}/\text{cm}^2$) | | | |
| --- | --- | --- | --- | --- | --- | --- | --- | --- | --- | --- | --- | --- |
|  | Experiment 1 |  | Experiment 2 |  | Experiment 1 |  | Experiment 2 |  | Experiment 1 |  | Experiment 2 |  |
|  | EMM (SE) | CV (%) | EMM (SE) | CV (%) | EMM (SE) | CV (%) | EMM (SE) | CV (%) | EMM (SE) | CV (%) | EMM (SE) | CV (%) |
| ARZM03042 | 46 (3.43)abc | 34 | 9.5 (0.35)abc | 46 | 8 (0.34)abc | 8.84 | 9.5 (0.35)a | 13.59 | 37.1 (3.12)ab | 34.2 | 35.1 (0.89)a | 33.7 |
| ARZM04013 | 37.2 (2.77)ab | 33 | 10.4 (0.33)abc | 51 | 7.6 (0.33)abc | 7.21 | 10.4 (0.33)ab | 8.6 | 35.7 (3)ab | 29 | 41.3 (0.62)b | 40 |
| ARZM04029 | 43.5 (3.24)abc | 33 | 11 (0.75)abc | 63 | 7.8 (0.34)abc | 10.73 | 11 (0.75)ab | 0 | 38.8 (3.26)ab | 34.6 | 37.4 (1.2)ab | 36.2 |
| ARZM04060 | 50.1 (4.17)abc | 45 | 10.7 (0.37)ab | 44 | 8.75 (0.40)abc | 5.71 | 10.7 (0.37)ab | 4.65 | 44 (3.7)b | 27.2 | 40 (3.31)ab | 33.2 |
| ARZM05007 | 44.4 (3.3)abc | 34 | 10.2 (0.32)abc | 45 | 8.2 (0.35)abc | 10.2 | 10.2 (0.32)ab | 4.38 | 33.3 (2.8)ab | 24.6 | 42.2 (2.82)ab | 34 |
| ARZM05067 | 49.8 (3.71)abc | 37 | 10.2 (0.32)abc | 51 | 8.4 (0.35)abc | 6.52 | 10.2 (0.32)ab | 4.38 | 26.6 (2.23)a | 20.2 | 37.6 (2.07)ab | 32 |
| ARZM05069 | 47.6 (3.54)abc | 38 | 10.4 (0.33)bc | 55 | 9.2 (0.36)c | 9.09 | 10.4 (0.33)ab | 5.27 | 36.5 (3.06)ab | 28.3 | 34.2 (1.39)a | 30.7 |
| ARZM05118 | 58.2 (4.34)c | 48 | 11.6 (0.34)bc | 55 | 8.4 (0.35)abc | 13.57 | 11.6 (0.34)b | 13.07 | 41.7 (3.5)b | 38.4 | 40.4 (2.16)ab | 33.5 |
| ARZM05120 | 44.9 (3.34)abc | 33 | 10.2 (0.32)abc | 48 | 7.4 (0.33)ab | 7.4 | 10.2 (0.32)ab | 10.74 | 45.4 (3.82)b | 40.7 | 40.6 (2.69)ab | 34.2 |
| ARZM06060 | 34.6 (2.57)a | 22 | 10 (0.36)abc | 45 | 7 (0.32)a | 10.1 | 10 (0.36)ab | 8.16 | 35.2 (2.96)ab | 27.3 | 41.9 (2.01)ab | 36.6 |
| ARZM06109 | 46.7 (3.48)abc | 39 | 10.3 (0.36)a | 39 | 8.2 (0.35)abc | 13.36 | 10.3 (0.36)ab | 4.88 | 47.4 (3.99)b | 41 | 40.3 (2.17)ab | 33.3 |
| ARZM08018 | 48.6 (4.05)abc | 44 | 10.8 (0.33)bc | 60 | 9 (0.40)bc | 18.14 | 10.8 (0.33)ab | 7.75 | 42.3 (3.55)b | 37.3 | 40.9 (1.7)ab | 34.7 |
| ARZM08020 | 41.7 (3.11)abc | 33 | 10 (0.32)abc | 32 | 8.6 (0.35)abc | 13.26 | 10 (0.32)ab | 7.07 | 42.7 (4.02)b | 35.6 | 42.5 (2.53)ab | 35.7 |
| ARZM08096 | 53.2 (3.96)bc | 49 | 11 (0.33)abc | 55 | 9 (0.36)bc | 7.86 | 11 (0.33)ab | 11.13 | 34 (2.86)ab | 23.2 | 40 (1.73)ab | 34 |
| ARZM08178 | 53.2 (3.96)bc | 48.5 | 10.6 (0.33)c | 54 | 9 (0.36)bc | 7.86 | 10.6 (0.33)ab | 5.17 | 38.2 (3.21)ab | 33.5 | 40.7 (4.25)ab | 30.2 |
| ARZM10036 | 45.6 (3.4)abc | 39.5 | 10.4 (0.33)bc | 60 | 9 (0.36)bc | 13.61 | 10.4 (0.33)ab | 5.27 | 39.5 (3.32)ab | 34 | 39.2 (2.654)ab | 33.3 |
| ARZM10076 | 50.8 (3.78)bc | 43 | 11 (0.33)abc | 49 | 8.8 (0.36)bc | 9.51 | 11 (0.33)ab | 6.43 | 43.8 (3.68)b | 37.6 | 41.2 (3.26)ab | 31.1 |
| ARZM10082 | 50 (3.73)abc | 47 | 10.6 (0.33)bc | 61 | 8.2 (0.35)abc | 10.2 | 10.6 (0.33)ab | 5.17 | 35.9 (3.02)ab | 25.8 | 41.2 (1.62)ab | 35.5 |
| ARZM12205 | 56.5 (4.71)c | 44 | 10 (0.32)bc | 59 | 8.75 (0.40)abc | 5.71 | 10 (0.32)ab | 7.07 | 34.7 (2.91)ab | 23.5 | 42.1 (1.37)b | 39.3 |
| Range | 34.6–58.2 |  | 9.5–11.6 |  | 7–9.2 |  | 9.5–11.6 |  | 26.6–47.4 |  | 34.2–42.5 |  |
| Maximum fold-change | 1.68 |  | 1.22 |  | 1.31 |  | 1.22 |  | 1.78 |  | 1.24 |  |

**Table 2.** Biomass and Stem Diameter in combined Experiments 1 and 2. Data represent Estimated Marginal Means or EMM (Standard Errors or SE) from Linear Mixed Effects Models, with different letters indicating significant differences (p<0.05), and Coefficients of Variation (CV). N=3-5.

| Accession | Total Biomass (g) |  | Leaf Biomass (g) |  | Stem Biomass (g) |  | Stem Diameter (cm) |  |
| --- | --- | --- | --- | --- | --- | --- | --- | --- |
|  | EMM (SE) | CV (%) | EMM (SE) | CV (%) | EMM (SE) | CV (%) | EMM (SE) | CV (%) |
| ARZM03042 | 42.8 (23.8)abc | 16.7 | 17.1 (11.22)ab | 7.25 | 15.43 (8.16)abcde | 5.65 | 1.05 (0.61)abc | 0.6 |
| ARZM04013 | 46.6 (25.5)abc | 7.28 | 21 (13.72)abc | 3.55 | 14.59 (7.66)abcde | 2.38 | 1.22 (0.71)c | 0.35 |
| ARZM04029 | 72.8 (40.2)bcd | 37.09 | 32.8 (21.69)bcd | 14.25 | 25.97 (14.08)cdef | 12.39 | 1.28 (0.75)c | 0.4 |
| ARZM04060 | 86.5 (47)cd | 42.58 | 33 (21.6)bcd | 13.37 | 27.35 (14.04)ef | 12.1 | 1.16 (0.68)abc | 0.42 |
| ARZM05007 | 73.2 (39.7)bcd | 20.37 | 24.6 (16.07)abcd | 9.03 | 19.8 (10.05)de | 7.31 | 1.10 (0.64)abc | 0.24 |
| ARZM05067 | 52.2 (28.5)abcd | 11.58 | 21.8 (14.23)abcd | 5.85 | 19.52 (10.09)bcdef | 5.36 | 1.06 (0.62)abc | 0.32 |
| ARZM05069 | 53.7 (29.3)abcd | 21.38 | 22.7 (14.83)abcd | 9.34 | 18.15 (9.21)cde | 5.54 | 1.10 (0.64)abc | 0.32 |
| ARZM05118 | 103.7 (56)d | 32.08 | 50.4 (32.89)d | 10.43 | 31.57 (16.12)f | 15.42 | 1.26 (0.73)c | 0.38 |
| ARZM05120 | 71 (38.5)bcd | 18.42 | 31.7 (20.69)bcd | 6.08 | 17.07 (8.68)bcd | 7.78 | 1.11 (0.65)abc | 0.53 |
| ARZM06060 | 26.9 (14.9)a | 5.44 | 11.1 (7.24)a | 2.43 | 8.77 (4.51)a | 3.22 | 0.86 (0.50)a | 0.22 |
| ARZM06109 | 48.2 (26.3)abc | 13.37 | 16.2 (10.59)ab | 6.91 | 14.38 (7.28)bc | 3.6 | 1.05 (0.61)abc | 0.3 |
| ARZM08018 | 83 (44.9)cd | 29.13 | 44.8 (29.23)cd | 8.68 | 25.97 (13.27)ef | 15.91 | 1.30 (0.76)c | 0.68 |
| ARZM08020 | 35.6 (19.7)ab | 7.35 | 15.5 (10.13)ab | 4.24 | 11.38 (5.85)ab | 3.05 | 0.86 (0.50)ab | 0.2 |
| ARZM08096 | 68 (36.8)bcd | 21.85 | 32.4 (21.04)bcd | 9.9 | 25.66 (13.34)def | 7.39 | 1.24 (0.72)c | 0.62 |
| ARZM08178 | 72.7 (39.9)bcd | 12.57 | 34.6 (22.77)bcd | 3.22 | 20.94 (11.66)abcdef | 3.87 | 1.31 (0.77)c | 0.3 |
| ARZM10036 | 65.5 (36)bcd | 27.3 | 29.7 (19.55)bcd | 9.4 | 19.41 (9.82)de | 10.15 | 1.16 (0.68)abc | 0.74 |
| ARZM10076 | 85.1 (45.9)cd | 35.53 | 45.6 (29.66)cd | 10.95 | 27.18 (14.31)def | 14.18 | 1.15 (0.67)bc | 0.65 |
| ARZM10082 | 64.1 (34.8)bcd | 14.32 | 23 (14.99)abcd | 5.93 | 15.8 (8.12)bcd | 7.03 | 1.25 (0.73)c | 0.36 |
| ARZM12205 | 62.5 (34)bcd | 21.76 | 32.3 (21.05)bcd | 7.74 | 17.27 (8.92)bcde | 8.8 | 1.10 (0.64)abc | 0.34 |
| Range | 26.9–103.7 |  | 11.1–50.4 |  | 8.77–31.57 |  | 0.86–1.31 |  |
| Maximum fold-change | 3.86 |  | 4.54 |  | 3.6 |  | 1.52 |  |

The analysis of phenotypic traits between accessions from NEA and NWA showed slight differences when compared at the region of origin level, with only two traits exhibiting statistically significant variation (Supplementary table 2). Leaf number differed significantly in Experiment 1, with NWA accessions producing more leaves (8.79) than NEA accessions (8.07), while NWA accessions had significantly greater plant height (64.1 cm) compared to those from NEA (56.8 cm) in Experiment 2. Other measured parameters, such as total chlorophyll content (measured with Dualex®), total biomass, leaf biomass, stem biomass, and stem diameter, showed no significant differences at the Region level across both experiments (p-values > 0.05). These results suggest that while some differences in plant height and leaf number were observed between the regions, most measured phenotypic traits remained largely unaffected by regional origin.

Next, variables were analyzed at the Accession level (Tables 1 and 2). Height marginal means ranged from 34.6 to 58.2 cm (average CV 39%) in Experiment 1 and 9.5 to 11.6 cm (average CV 51%) in Experiment 2 (Table 1), with three statistical groups in each experiment. ARZM05118 exhibited the greatest mean height in both experiments, while ARZM06060 and ARZM03042 had the lowest mean heights in Experiments 1 and 2, respectively. The number of leaves ranged from 7 to 9.2 in Experiment 1 and from 9.5 to 11.6 in Experiment 2, with three and two statistical groups in each experiment, respectively (Table 1). Both experiments showed small coefficients of variation (<10%). Total chlorophyll (Dualex®) ranged from 26.6 to 47.4 μg/cm² in Experiment 1 and 34.2 to 42.5 μg/cm² in Experiment 2 (Table 1). In both experiments, chlorophyll values differed moderately among accessions, with two statistical groups each, and CV <35%. The maximum fold-changes between minimum and maximum values varied between 1.22 and 1.78 for each trait (Table 1).

Total biomass ranged from 26.9 g (ARZM06060) to 103.7 g (ARZM05118) with four statistical groups, although most accessions clustered within the intermediate abc-bcd groups, while CV varied between 5-43% (Table 2). Leaf biomass ranged from 11.1-50.4g (CV < 14%, 4 statistical groups) while stem biomass ranged from 8.77-31.57g (CV < 16%, 6 statistical groups) (Table 2). Consistently with total biomass, ARZM06060 presented the lowest values and ARZM05118 the highest. Stem diameter varied less across landraces, ranging from 0.86 cm (ARZM06060) to 1.31 cm (ARZM08178), with generally low coefficients of variation (<0.74%) and three statistical groups (Table 2). Overall, the ranking of accessions was comparable across traits: high-biomass landraces (e.g., ARZM05118, ARZM10076, ARZM08118, ARZM08018) also produced greater leaf and stem biomass and tended to have larger stem diameters, while low-biomass accessions (e.g., ARZM06060, ARZM08020, ARZM03042) were consistently grouped among the lowest performers. This can be explained by the high correlation between these traits (Supplementary figure 2). On average, coefficients of variation were low to moderate for all traits, except for total biomass, indicating low variation within accessions. However, the fold-change between the maximum and minimum values for each trait was consistently high (range: 1.52-1.78), indicating substantial variation among the accessions.

Overall, although no differences were generally observed at the Region level between NWA and NEA groups (Supplementary table 2), morphological analysis showed variability among accessions with several significantly different groups for each trait (Tables 1 and 2). Although some accessions showed high variability within the accession (such as height in Experiment 2, average CV: 51.16%), most traits showed moderate or low coefficients of variation (<30% on average). It is worth highlighting that for all traits significant Accession effects were observed (p < 0.05), indicating a genetic influence linked to the accessions.

### Accessions Show Large Fold-Changes and High Within-Group Variation in Biochemical Variables

The metabolic content of plants influences several physiological processes with agronomical impact (Pott et al. 2019). We measured leaf metabolites involved in primary metabolism that affects plant growth and yield (total protein, total soluble sugars, starch, pigments) and secondary metabolites involved in general stress responses (total phenolics, total antioxidant activity). In coincidence with what was observed for morphological traits, biochemical traits showed higher CV in Experiment 1 than in Experiment 2, except for starch content and antioxidant activity (Supplementary figure 1).

The biochemical analysis of accessions from NEA and NWA revealed some regional trends in various traits, though none of them were statistically significant (Supplementary table 3). Accessions from NWA generally exhibited higher total proteins (1.52 mg/g), total soluble sugars (13.6 mg/g) and total carotenoids (0.957 g/g) compared to those from NEA (1.45 mg/g, 11.2 mg/g and 0.46 g/g, respectively). In contrast, NEA accessions had higher starch (11.79 mg/g), total chlorophyll (1.943 g/g) and chlorophyll a (1.55 g/g) compared to NWA accessions (8.53 mg/g, 0.987 g/g and 0.845 g/g, respectively). Chlorophyll b and total carotenoid levels were greater in NWA accessions (0.914 g/g and 0.957 g/g) than in NEA accessions (0.495 g/g and 0.46 g/g). Total phenolics and antioxidant activity (in both hydrophilic and hydrophobic phases) showed minimal regional variation. Beyond these trends, the variation within each region was considerable (CV 25-105%), reflecting a high degree of heterogeneity among accessions. These findings suggest that while regional differences may exist in certain biochemical traits, variability within regions may play a significant role in shaping these traits.

Next, variables were compared at the Accession level (Tables 3a and b). Total proteins ranged from approximately 2.2 (ARZM05067) to 7.3 mg/g (ARZM04013), with low coefficients of variation and two statistical clusters (Table 3a). Total soluble sugars varied more widely, from 7.7 (ARZM03042) to 16.2 mg/g (ARZM05007), while all landraces belonged to a single statistical group (Table 3a). Starch content showed high variability (CV frequently > 70%), with means spanning 6.7 (ARZM08020) to 17.9 mg/g (ARZM06109), while no significant differences were observed among accessions (Table 3a). Total phenols ranged from 1.86 (ARZM10076) to 3.03 mg GA/g (ARZM08020), forming two statistical groups with CV lower than 1.84% (Table 3a). Both measures of antioxidant activity (hydrophilic and hydrophobic phases) showed very limited differentiation among accessions (Table 3a). Antioxidant activity in hydrophilic phase values ranged from 0.076 (ARZM04029) to 0.104 mg GA/g (ARZM08020), while antioxidant activity in hydrophobic phase ranged from 0.053 (ARZM08020) to 0.146 mg GA/g (ARZM12205), all within a single significance group, with high CV for hydrophilic phase (>46%) and low CV for the hydrophobic phase measurements (<0.1%). In contrast, maximum fold-changes when comparing maximum and minimum values for each trait were high (range: 1.37-3.33), indicating high variability between accessions.

**Table 3a.** Leaf metabolite content of fully expanded leaves from V12 stage plants in combined Experiments 1 and 2 (Part 1). Data represent Estimated Marginal Means or EMM (Standard Errors or SE) from Linear Mixed Effects Models, with different letters indicating significant differences (p<0.05), and Coefficients of Variation (CV). N=3-5.

| Accessions | Total Proteins (mg/g) |  | Total Soluble Sugars (mg/g) |  | Starch (mg/g) |  | Total Phenols (mg GA/g) |  | Antioxidant Activity (Hydrophilic phase) (mg GA/g) |  | Antioxidant Activity (Hydrophobic phase) (mg GA/g) |  |
| --- | --- | --- | --- | --- | --- | --- | --- | --- | --- | --- | --- | --- |
|  | EMM (SE) | CV (%) | EMM (SE) | CV (%) | EMM (SE) | CV (%) | EMM (SE) | CV (%) | EMM (SE) | CV (%) | EMM (SE) | CV (%) |
| ARZM03042 | 6.33 (2.3)ab | 4.34 | 7.73 (1.49)a | 5.21 | 8.6 (10.12)a | 113.2 | 1.94 (0.28)ab | 1.2 | 0.0967 (0.0607)a | 52.41 | 0.078 (0.0219)a | 0.03 |
| ARZM04013 | 7.27 (2.61)b | 3.17 | 11.43 (2.2)a | 5.9 | 11.43 (13.37)a | 56.71 | 1.86 (0.26)a | 1.17 | 0.0924 (0.0604)a | 65.86 | 0.1026 (0.0263)a | 0.06 |
| ARZM04029 | 3.5 (1.29)a | 2.86 | 9.63 (2.14)a | 6 | 11.72 (13.66)a | 65.8 | 1.89 (0.29)ab | 1.11 | 0.076 (0.0624)a | 60.52 | 0.0604 (0.0189)a | 0.03 |
| ARZM04060 | 5.92 (2.29)ab | 2.07 | 9.63 (1.85)a | 4.03 | 10.99 (12.86)a | 72.82 | 2.08 (0.28)ab | 1.36 | 0.0913 (0.0605)a | 57.83 | 0.0682 (0.0194)a | 0.004 |
| ARZM05007 | 3.09 (1.25)a | 1.1 | 16.23 (2.79)a | 5.06 | 17.73 (20.94)a | 76.5 | 2.18 (0.25)ab | 1.54 | 0.085 (0.0607)a | 64.24 | 0.0965 (0.0235)a | 0.03 |
| ARZM05067 | 2.18 (1.41)ab | 0.05 | 10.49 (1.9)a | 7.37 | 8.44 (9.78)a | 120.5 | 2.1 (0.26)ab | 1.26 | 0.1017 (0.0612)a | 46.55 | 0.1207 (0.0395)a | 0.1 |
| ARZM05069 | 3.93 (1.51)ab | 2.18 | 13.9 (2.39)a | 4.54 | 11.42 (13.26)a | 77.1 | 2.33 (0.26)ab | 1.28 | 0.092 (0.0605)a | 75.49 | 0.0921 (0.0249)a | 0.01 |
| ARZM05118 | 2.73 (1.15)a | 0.37 | 8.16 (1.48)a | 2.93 | 11.36 (13.19)a | 72.55 | 1.92 (0.25)ab | 1.04 | 0.0932 (0.0605)a | 63.7 | 0.0859 (0.023)a | 0.03 |
| ARZM05120 | 6.63 (2.44)ab | 3.26 | 13.36 (2.57)a | 7.47 | 10.18 (11.88)a | 104.75 | 2.06 (0.25)ab | 1.65 | 0.0903 (0.0605)a | 72.07 | 0.0824 (0.021)a | 0.01 |
| ARZM06060 | 5.89 (2.15)ab | 2.89 | 12.45 (2.26)a | 6.02 | 12.65 (14.72)a | 103.29 | 2.34 (0.26)ab | 1.6 | 0.0989 (0.0605)a | 63.48 | 0.1124 (0.0269)a | 0.02 |
| ARZM06109 | 3.12 (1.26)a | 1.16 | 11.8 (2.03)a | 6.8 | 17.9 (20.98)a | 94.44 | 2.03 (0.26)ab | 0.87 | 0.0997 (0.0606)a | 61.28 | 0.0952 (0.0232)a | 0.03 |
| ARZM08018 | 5.57 (2.07)ab | 3.2 | 14.75 (2.54)a | 4.83 | 8.28 (9.66)a | 137.59 | 2.7 (0.25)ab | 1.84 | 0.0907 (0.0605)a | 72.59 | 0.0984 (0.0258)a | 0.03 |
| ARZM08020 | 3.44 (1.36)ab | 1.91 | 14.22 (3.16)a | 4.5 | 6.74 (8.24)a | 166.26 | 3.03 (0.27)b | 1.49 | 0.1044 (0.0612)a | 54.29 | 0.0533 (0.0174)a | 0.01 |
| ARZM08096 | 4.67 (1.68)ab | 2.11 | 11.75 (2.13)a | 6.37 | 8.13 (9.42)a | 90.02 | 2.14 (0.25)ab | 1.34 | 0.0861 (0.0605)a | 78.65 | 0.0922 (0.0282)a | 0.01 |
| ARZM08178 | 3.41 (1.5)ab | 0.71 | 13.81 (2.66)a | 8.3 | 9.37 (10.94)a | 104.5 | 2.24 (0.26)ab | 1.5 | 0.0854 (0.0605)a | 67.96 | 0.1055 (0.0271)a | 0.03 |
| ARZM10036 | 5.2 (1.87)ab | 3.46 | 13.09 (2.37)a | 1.39 | 8.99 (10.47)a | 101.48 | 1.98 (0.25)ab | 1.33 | 0.0888 (0.0605)a | 70.5 | 0.0898 (0.0217)a | 0.02 |
| ARZM10076 | 5.27 (1.97)ab | 2.68 | 13.42 (2.58)a | 5.87 | 13.84 (16.14)a | 88.04 | 1.84 (0.26)a | 1.31 | 0.0925 (0.0606)a | 77.23 | 0.1087 (0.0298)a | 0.04 |
| ARZM10082 | 5.23 (1.94)ab | 2.69 | 13.37 (2.42)a | 7.12 | 7.21 (8.52)a | 108.31 | 2.12 (0.25)ab | 1.51 | 0.0951 (0.0605)a | 67.26 | 0.0806 (0.0202)a | 0.01 |
| ARZM12205 | 4.3 (1.76)ab | 0.89 | 14.61 (2.65)a | 8.27 | 8.78 (10.22)a | 85.74 | 2.1 (0.25)ab | 1.53 | 0.0919 (0.0619)a | 59.03 | 0.1459 (0.0393)a | 0.01 |
| Range | 2.18–7.27 |  | 7.73–16.23 |  | 6.74–17.9 |  | 1.84–3.03 |  | 0.08–0.1 |  | 0.05–0.15 |  |
| Maximum fold-change | 3.33 |  | 2.1 |  | 2.66 |  | 1.65 |  | 1.37 |  | 2.74 |  |

GA= gallic acid.

Total chlorophyll ranged from 1.01 to 2.68 g/g, with ARZM05120 displaying the highest mean value, forming two statistical clusters (Table 3b). Several accessions -including ARZM08020, ARZM08178, and ARZM12205, and ARZM05067-showed the lowest total chlorophyll levels. For chlorophyll a, values ranged from 0.72 (ARZM08020 and ARZM12205) to 2.48 g/g (ARZM05120), while chlorophyll b ranged from 0.72 (ARZM05067) to 1.23 g/g (ARZM10082) (Table 3b). Chlorophyll a had three statistical groups. On the contrary, accessions did not show statistically significant differences for chlorophyll b. Total carotenoids showed a narrower range -between 0.86 (ARZM05007) and 1.26 g/g (ARZM05118)-, while accessions did not show statistical differences (Table 3b). All the chlorophylls as well as the carotenoids showed high CV values (24-107%), suggesting substantial within-accession variability. Maximum fold-changes when comparing maximum and minimum values for each trait were high (range: 1.45-3.46), indicating high variability between accessions. Chlorophylls were highly correlated with each other (Supplementary figure 2). In turn, they were positively correlated with growth-related parameters (biomass, height, number of leaves), possibly due to their relationship with photosynthesis.

**Table 3b.**
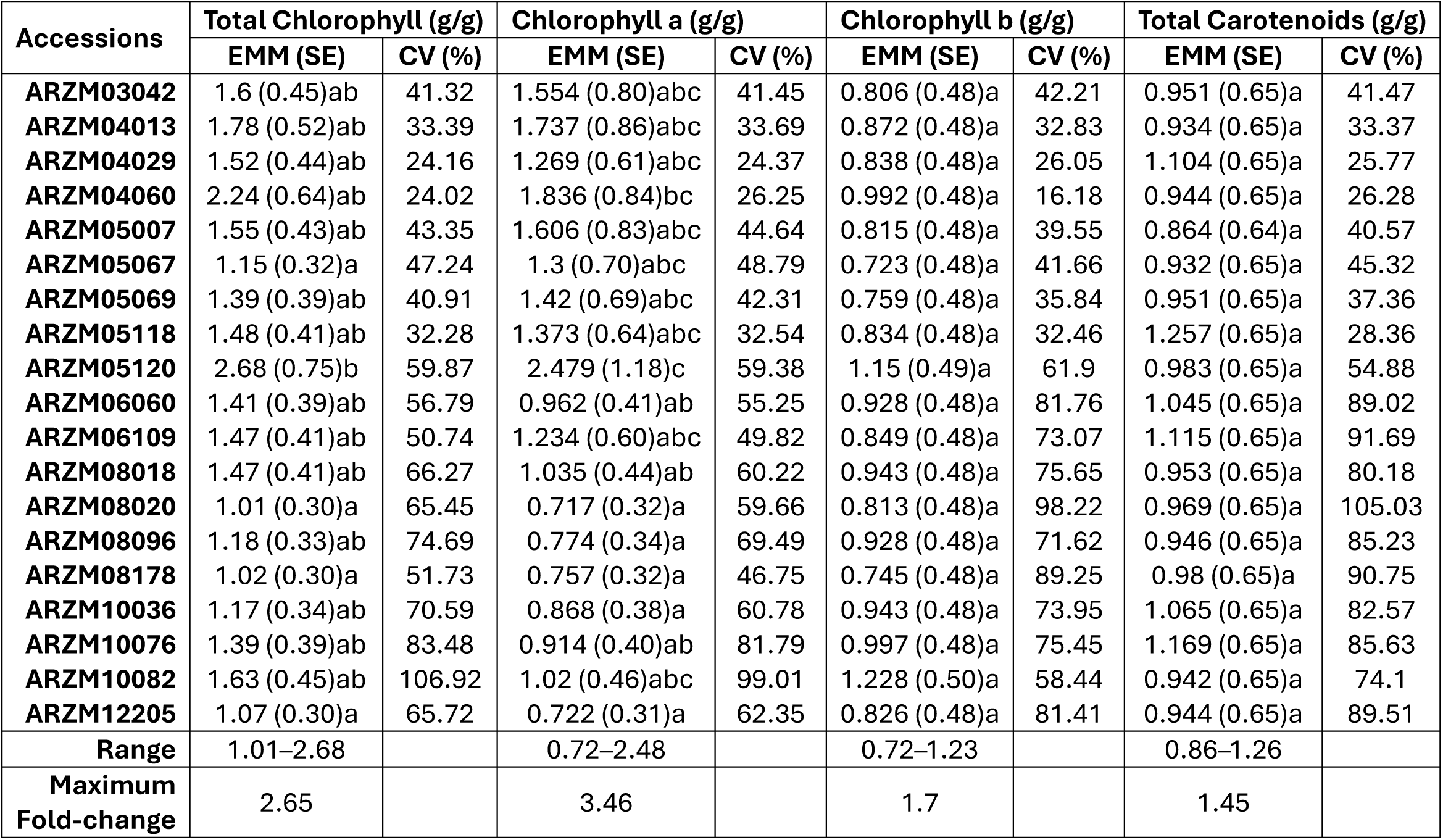
Leaf metabolite content of fully expanded leaves from V12 stage plants in combined Experiments 1 and 2 (Part 2). Data represent Estimated Marginal Means or EMM (Standard Errors or SE) from Linear Mixed Effects Models, with different letters indicating significant differences (p<0.05), and Coefficients of Variation (CV). N=3-5.

Total proteins, total phenols, total chlorophyll, and chlorophyll a exhibited significant Accession effects (p < 0.05), indicating a genetic influence linked to the accessions. While the analysis of biochemical traits did not reveal many statistical groups or clusters -likely due to the limited sample size per accession-there was considerable variability in each trait observed both between and within accessions. Between accessions, the maximum value was as high as ≈3.5-fold higher than the minimum (for chlorophyll a and total proteins) and as low as ≈1.5-fold higher than the minimum (for total carotenoids and antioxidant activity in hydrophilic phase) (Table 3a and b). Additionally, significant variation was seen within each accession, as evidenced by the high CV in some accessions, with starch showing the highest average CV of 96% (Table 3a). This variability between and within accessions might hold biological significance.

### Salt Stress Tolerance Varies by Accessions and Regions

With the aim of extending the phenotypic characterization of the 19 accessions, we assessed the performance of their seedlings to salt stress, considering that the likelihood of soil salinity increases with climate change (Corwin 2021), affecting many crop species including maize (Farooq et al. 2015; Hu et al. 2022). When compared at the Region level, the Region × Treatment interaction was non-significant, so Region was averaged across Treatment (NaCl concentration). NWA plants showed higher root weight and length in comparison to NEA plants, while no differences were observed at the leaf level (Figure 2a and b). This suggests that there is a difference in the response of seedlings according to their region of origin.

**Figure 2.**
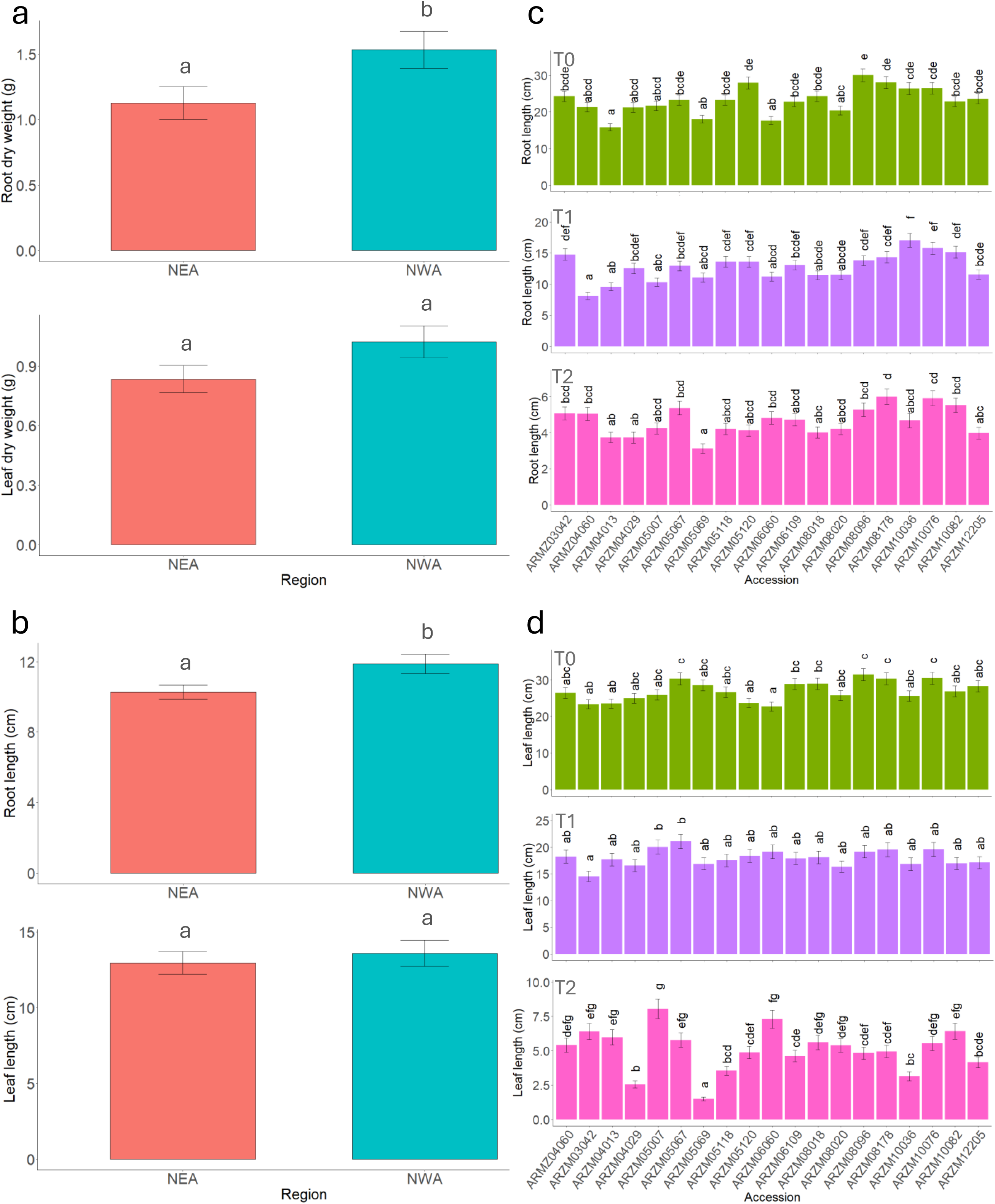
Salt stress tolerance assay. (a) Dry weight of roots (upper panel) and leaves (lower panel) by Region. (b) Length of roots (upper panel) and leaves (lower panel) by Region. Length of roots (c) and leaves (d) by Accession. Data represent Estimated Marginal Means or EMM (Standard Errors or SE) from Linear Mixed Effects Models, with different letters indicating significant differences (p<0.05). Three independent repetitions were performed with N=19 (a,b) and N=20 (c,d). T0 (control, 0 mM NaCl), T1 (175 mM NaCl), T2 (350 mM NaCl). NEA: Northeastern Argentina. NWA: Northwestern Argentina.

When compared at the Accession level, Accession × Treatment interaction was significant for both root and leaf length; thus, the effect of each level of Treatment was evaluated separately across Accessions (Figure 2c and d). Overall, root length decreased over time from T0 to T2 across all accessions (Figure 2c). It showed some variation among accessions under control conditions (T0), with mean values ranging from 17.78 to 30.09 cm. Under the T1 and T2 treatments, differentiation among accessions was also evident, with mean values ranging from 8.11 to 17.06 cm (T1) and 3.12 to 6 cm (T2). In all cases, ARZM05069 and ARZM04029 showed consistently lower values, while ARZM10076, ARZM08178 and ARZM08096 showed higher values.

Leaf length also decreased from T0 to T2 across Accessions (Figure 2d). Under control conditions (T0), it showed variation among accessions, with mean values ranging from 22.68 to 31.44 cm. Under treatment T1, accession means ranged from 14.57 to 21.13 cm, with most genotypes not differing significantly. Under treatment T2, values were markedly lower, ranging from 1.49 cm in ARZM05069 to 8.07 cm in ARZM05007, with seven statistically significant groups. ARZM05067 appeared consistently among the highest means.

Altogether, both T1 and T2 treatments had a marked effect across accessions, significantly reducing the measured variables compared with control conditions (T0). Both T1 and T2 treatments revealed clear differences among accessions at both root and leaf levels, indicating genotype-specific responses. Across both measured variables (Figure 2c and d), ARZM05069 and ARZM04029 showed lower means at T2, indicating higher sensitivity, while ARZM05067 and ARZM10082 showed higher means at T2, indicating higher tolerance. When considering regional and accession differences, the root phenotype seems to be more responsive to the treatments than the leaf phenotype, indicating a more prominent role in the response to salt stress in these plants.

### Regional Differences at the Phenotypic Level Can Be explained by the Altitudinal Cline

To explore global differences among regions and accessions, we examined the relationship between morphological and biochemical traits measured in adult plants (Tables 1, 2, and 3) using a combination of multivariate analyses. The Principal Components Analysis (PCA) revealed high phenotypic variation among the different maize Accessions and different stability of these accessions across the two experiments (Figure 3a). Accessions with the highest overall variability, indicated by a wide dispersion of points and greater distance from their centroids included ARZM10076, ARZM05007 and ARZM05120 (Figure 3a). Conversely, populations with the lowest variability and highest stability were ARZM08018, ARZM05069 and ARZM08020, which demonstrated consistent performance between the two experiments (Figure 3a). Regarding explained variance and variables contribution, PC1 explained 31.40% of the variance, mainly influenced by biomass and pigments, while PC2 explained 22.80% of the variance, with pigments and biomass contributing most as well (Figure 3b). The PCA biplot shows that accessions are distributed along both axes, with no clear pattern of separation by experiment or region of origin (Figure 3a; Supplementary Figure 3). In turn, based on the BIC criterion, the DAPC identified k = 4 as the most probable number of clusters (Figure 3c, d), and eleven accessions were consistently grouped in both experiments, although no clustering by region was observed in agreement with the PCA. The grouping was highly concordant with the major variation axes identified by the PCA (Figure 3a, b). Accessions such as ARZM3042 in Experiment 2, and ARZM05120 and ARZM10082 in Experiment 1, were grouped together into Cluster 1 in the DAPC (Figure 3c, d) and had negative PC1 scores (Figure 3a, b), characterized by high-moderate values for pigments and moderate-low biomass values (Tables 2, 3b). A second group (Cluster 2 of the DAPC, Figure 3c and d) including ARZM04013, ARZM05067, ARZM05069, ARZM06109, and ARZM08020, were distinctively defined by moderate-low pigment values and low biomass values (Tables 2 and 3b) and had positive scores on PC2 of the PCA (Figure 3a, b). In contrast, accessions ARZM04029, ARZM05118, ARZM08096, and ARZM8178 were grouped in Cluster 3 of the DAPC (Figure 3c, d), had positive scores on PC1 of the PCA (Figure 3a, b), and were characterized by high-moderate biomass and moderate-low pigment values (Tables 2 and 3b). Finally, populations including ARZM08018 and ARZM10076 were grouped into Cluster 4 of the DAPC (Figure 3c, d) and had negative scores on the PC2 of the PCA (Figure 3a, b), based on their high biomass values, and moderate pigment content (Tables 2 and 3b).

**Figure 3.**
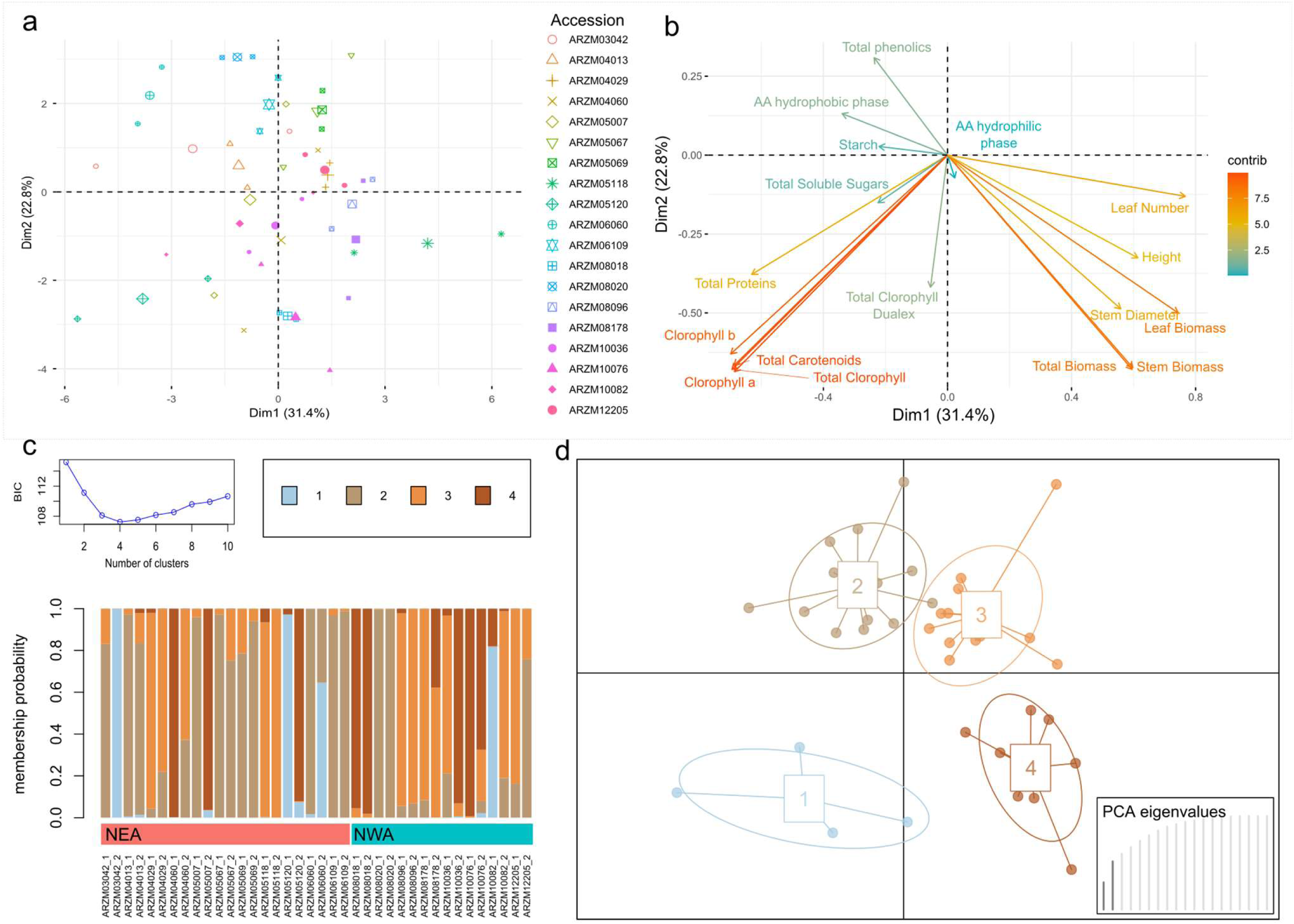
Multivariate ordination and clustering analyses. **a**. PCA biplot based on accession and experiment mean values. Each accession is represented by two smaller points corresponding to the means of Experiment 1 and 2. Larger points indicate centroids. **b.** Variable contributions to the first two principal components of the PCA. **c.** Membership probabilities of each accession in Experiments 1 and 2 for the four DAPC clusters (lower panel) selected according to BIC (upper panel). **d.** Scatterplot of the four DAPC clusters. AA: Antioxidant Activity. NWA: Northwestern Argentina. NEA: Northeastern Argentina.

Hierarchical cluster analysis, which was performed on marginal means from univariate linear models, identified five groups as the optimal, according to the average silhouette criterion (Figure 4a). Group 1 comprised accessions ARZM10076 and ARZM05118; Group 2 included ARZM04060, ARZM04013, and ARZM05120; Group 3 contained ARZM04029, ARZM03042, and ARZM05067; Group 4 consisted solely of ARZM08020; and Group 5 encompassed the remaining ten accessions. No evident and strong clustering pattern associated with the region of origin was observed. Pairwise distances highlighted that these groups display marked internal similarity and strong divergence from the others (Figure 4b). Group 4 (ARZM08020) was the most divergent, which showed high distance values relative to nearly all other accessions. Additionally, groups 1 and 2 also appeared highly distinct, exhibiting very low internal distances but strong divergence from the other accessions. In contrast, the larger group containing the remaining ten accessions (Group 5) displayed more moderate and mixed distance values, indicating some internal structure.

**Figure 4.**
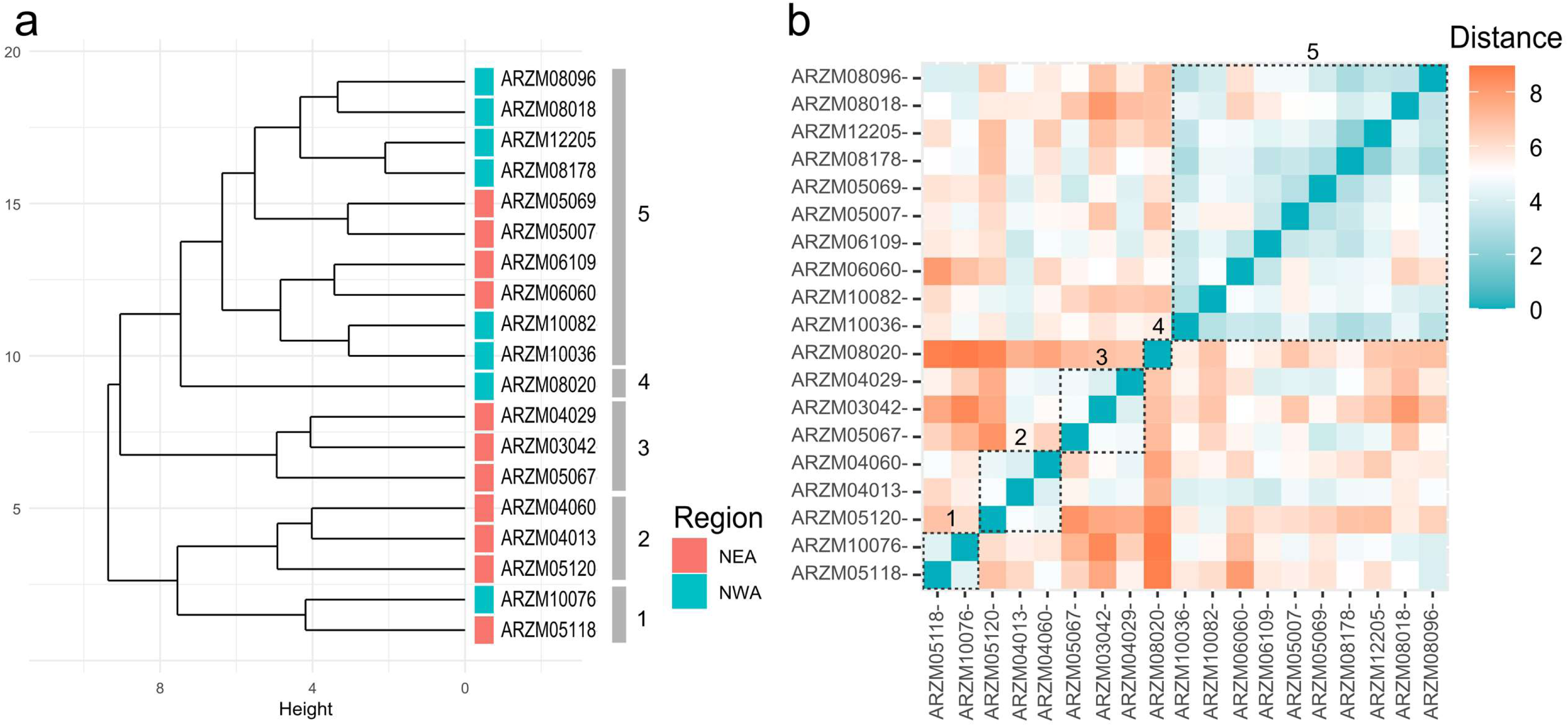
Hierarchical clustering. a. Dendrogram based on Euclidean distances, showing cluster assignment and region of origin of each accession. b. Heatmap of pairwise Euclidean distances among accessions. Dashed lines indicate the assigned cluster number. The clustering was done employing marginal means (Tables 1-3). NWA: Northwestern Argentina. NEA: Northeastern Argentina.

The clustering similarity between the dendrogram and the PCA/DAPC results was moderate, with a stronger alignment to the PCA outcomes (Figures 3, 4). Given that the clustering was based on marginal means from univariate linear models, whereas the PCA/DAPC analyses used raw means, this suggests that the observed differences between units at the multivariate level are, at least in part, influenced by their varying responses to the environment.

Redundancy Analysis (RDA) used for the association between multivariate phenotypic variation and geographic predictors revealed an altitudinal pattern. The first RDA model including Altitude as the sole predictor showed that elevation accounted for a significant portion of multivariate phenotypic variation (13.5%, *F* = 2.49, *p* = 0.02). This indicated that Altitude alone captures a detectable ecological gradient associated with trait differentiation among populations. The full model including Altitude, Latitude, and Longitude, explained ∼25.3% of total variance, and only Altitude explained a significant portion of the trait variation (sequential test: *F* = 2.53, *p* = 0.029), whereas Latitude and Longitude showed no significant effects (*p* > 0.29). Marginal tests revealed that the three variables capture overlapping spatial structure and cannot be statistically disentangled due to spatial autocorrelation and collinearity. The differentiation is associated with Altitude, but it is partly a consequence of the spatial structure (Latitude/Longitude). The ordination biplot reflected this pattern (Figure 5), with a strong axis of variation aligned with Altitude. Accessions occurring at higher elevations (ARZM8096, ARZM8018, ARZM10082, ARZM12205), all from NWA, were displaced toward the positive end of RDA1, showing a clear separation from NEA low-elevation populations, which cluster on the opposite side of the axis. Latitude and Longitude vectors had lower contribution relative to Altitude. High-altitude samples (positive RDA1 scores) were associated with higher concentrations of total sugars, total phenolics, carotenoids, and chlorophyll b (Supplementary table 4). In contrast, low-altitude samples (negative RDA1 scores) were characterized by higher levels of chlorophyll a, total chlorophyll, starch, and hydrophilic antioxidant capacity (Supplementary table 4). This aligns with tendencies in univariate analyses (Supplementary table 3).

**Figure 5.**
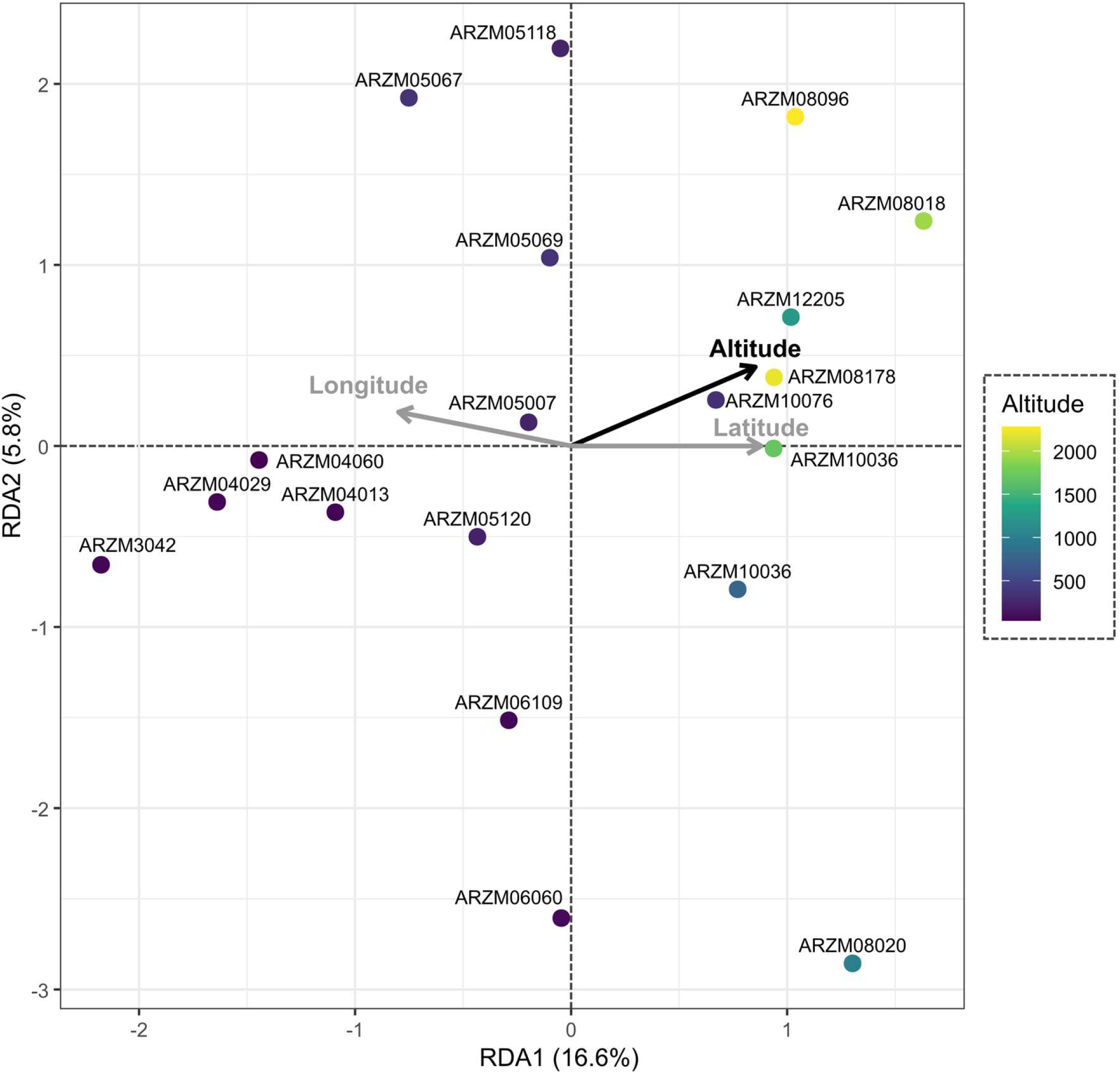
Redundancy analysis (RDA). Model biplot displaying the first two axes of the model including biochemical–morphological traits and the three environmental predictors (Elevation, Latitude, and Longitude). Elevation is highlighted in black, as it was the only significant predictor in the full model (p<0.05).

In conclusion, more than half of the accessions showed consistency in phenotypic characteristics across Experiments (Figure 3, 4), while the variables that contributed most to distinguishing the accessions were biomass and pigment content (Figure 3). Although the multivariate analyses did not reveal a clear regional separation (Figures 3c, 4a, and Supplementary figure 3), the RDA, which examined the contribution of environmental variables (Figure 5), indicated that altitude significantly influenced the phenotypic characteristics of the accessions. Therefore, it can be concluded that phenotypic variation among accessions followed an altitudinal gradient, driven largely by differences in pigment composition.

## Discussion

Studying population diversity is essential for the conservation and sustainable use of crop germplasm. In this work, we analyzed the morphological, biochemical, and salinity-stress tolerance variability of 19 native maize accessions from Northwestern and Northeastern Argentina, one of the southernmost regions in the world where maize landraces are cultivated, with the aim of providing information on agronomically relevant traits for maize breeding and conservation programs.

### Maize Landrace Accessions as a Source of Phenotypic Variation

Both univariate and multivariate statistical analyses revealed high variability among the 19 accessions evaluated in terms of morphology, biochemistry, and salt stress tolerance (Tables 1-3, Figures 2-4). The phenotypic variability observed is likely a reflection of the high genetic variability present in native maize varieties from Argentina (Bracco et al. 2016; Rivas et al. 2022; Dominguez et al. 2024a) and the Americas more broadly (Hufford et al. 2012). The origin of this extensive genetic variability in maize landraces can be attributed to multiple biocultural factors, including germplasm exchange among farmers, crosses between different local varieties due to their open-pollinated nature, farmer selection to meet quality and diversity requirements, adaptation to diverse climatic conditions, and gene flow with wild relatives (teosintes) in Mesoamerica (Camacho Villa et al. 2005; Arteaga et al. 2016; Guzzon et al. 2021).

The fact that the univariate analyses conducted in this study showed the accession factor to be statistically significant for all morphological variables and for several biochemical variables further supports the idea that the observed phenotypic variability has a genetic basis associated with each accession. This finding highlights the potential use of these accessions in crop improvement, which is particularly relevant given that commercial varieties harbor only between 5 and 10% of the available genetic variability in maize (Hoisington et al. 1999). In this regard, numerous examples underscore the importance of increasing the genetic diversity in commercial maize, such as the impact of fall armyworm (FAW; *Spodoptera frugiperda*) following the development of resistance to insecticides in some maize lines (Blanco et al. 2016), as well as the severe impact of drought on maize production in the United States in 2012 (Boyer et al. 2013). Nevertheless, despite their potential as sources of novel genetic variation and agronomically valuable traits, maize landraces have not been widely incorporated into breeding programs (Dominguez et al. 2024b). Maize landraces from other regions of the Americas have been used to improve yield under drought conditions (Medeiros Barbosa et al. 2021) and increase resistance to the tar spot complex (Willcox et al. 2022). Together, these studies highlight the importance of introducing the diversity present in landraces into breeding populations and emphasize the need to characterize the phenotypic variability of native maize as a resource for crop improvement.

Many of the variables measured in this study, for which differences among accessions were detected both statistically and in terms of magnitude as measured by fold-changes, have high agronomic relevance and could be of interest for breeding programs. At the phenotypic level, a larger stem diameter may be desirable for increasing water and nutrient transport capacity or for providing greater mechanical resistance, thereby reducing the risk of lodging (García et al. 2001). Leaf number and biomass are traits associated with the photosynthetic capacity of maize (Mehta and Sarkar 1992) and its potential use as forage or for bioenergy production (Klopfenstein et al. 2013). At the biochemical level, phenolic compounds play important roles in responses to both biotic and abiotic stress (Tak and Kumar 2021). Sugars, proteins, and starch influence leaf quality in maize used as a forage species (Liimatainen et al. 2022), while starch also plays a key role in seed germination and early seedling growth until photosynthetic maturity is reached, among other functions (Li et al. 2025). In addition, salinity tolerance has important implications for food security, not only in regions with saline soils (Egea et al. 2023) but also in relation to other stress conditions, given the extensive crosstalk between stress signaling pathways (Mishra et al. 2016; Kesawat et al. 2023).

Overall, the accessions exhibited high phenotypic variability across nearly all measured variables, with each accession displaying a unique combination of traits, as well as substantial variability within accessions (Tables 1-3). These characteristics demonstrate the considerable potential of these accessions as a source of phenotypic variation of agronomic interest and suggest that these populations may be capable of responding to a wide range of environmental pressures and future challenges.

### Altitudinal Cline as a Key Variable Driving Differences Between Regions

Although no significant differences were detected when variables were compared at the regional level (NWA vs. NEA) (Supplementary Tables 2 and 3), the RDA indicated that multivariate phenotypic variability among accessions could be explained by altitude (Figure 5). All accessions from the NEA region originate from altitudes below 345 m.a.s.l., whereas all accessions from the NWA region originate from altitudes above 782 m.a.s.l., with one exception (accession ARZM10076, originating at 322 m.a.s.l.) (Supplementary Table 1). Therefore, the phenotypic differentiation between regions appears to be driven primarily by altitude. RDA may have been more effective than linear mixed models in detecting these phenotypic differences because it captures the gradual altitudinal cline, whereas the linear mixed models may have overlooked this more subtle environmental effect by treating region as a categorical variable.

Phenotypic differences, both morphological and biochemical, within and among species as a function of altitude have long been documented (Körner 1989; Terashima et al. 1995; Sundqvist et al. 2013; Ratier Backes et al. 2022). In maize, numerous studies report phenotypic differences -most often morphological-associated with altitude in Mexican (Pace et al. 2024), African (Asare et al. 2016), Brazilian (Ribeiro e Souza et al. 2008), and North and South American landraces more broadly (Romero-Navarro et al. 2017; López-Valdivia et al. 2025). A comprehensive overview of these studies is provided in Salve et al. (2023a). Notably, Janzen et al. (2022) identified altitude-associated phenotypic traits, such as increased anthocyanin content in highland maize from Mexico, as evidence of local adaptation, suggesting that altitudinally structured phenotypic variation may contribute to adaptive processes.

In Argentina, fewer studies have addressed altitude-related variation in maize, but existing work has reported genetic (Rivas et al. 2022), morphological (Salve et al. 2023b) and cytological (Realini et al. 2018) differences along altitudinal gradients. In our study, the RDA showed that several phenotypical traits contributed to the differentiation of the accessions including pigments and antioxidants. Because altitude-related phenotypic differences are driven by factors such as temperature, precipitation, solar irradiation, and other environmental variables (Sundqvist et al. 2013), a more detailed analysis of the relationships between phenotypic traits and specific environmental factors may reveal additional, previously undetected patterns in these populations.

### Maize Landraces as a Source of Salinity Stress Tolerance

Nearly half of the NWA individuals were collected from sites with Entisol soils, which are characterized by salinity in the upper 50 cm, while the remaining accessions originated from Mollisol soils, which can become saline under certain environmental conditions (Lavado 2007) (Supplementary Table 1). The observed association between regions of origin and greater tolerance to salt stress (Figure 2a and b) suggests that geographic origin may be linked to a process of local adaptation. With respect to their phenotype, NWA accessions exhibited longer and heavier roots (Figure 2a and b), a pattern consistent with previous reports showing that salt-tolerant maize often develops more extensive root systems (Farooq et al. 2015; Hu et al. 2022).

Pratt et al. (2022) evaluated 11 maize landraces from the Southwestern United States originating from regions with saline soils and identified one landrace that maintained high yield under saline field conditions. Together with our results, this finding suggests that native maize landraces represent a promising source of salinity stress tolerance and that further evaluation under field conditions, including comparisons with commercial lines, is warranted. In addition, tolerance to saline stress is known to exhibit substantial crosstalk with other stress-response pathways through shared signaling mechanisms, particularly hormonal pathways (such as abscisic acid signaling) and antioxidant responses, which can confer resistance to multiple stressors (Mishra et al. 2016; Kesawat et al. 2023). This interaction suggests the presence of additional stress tolerances that could be evaluated in future studies using these same accessions. For example, previous studies show that maize landraces from Argentina may offer a source of disease resistance (Defacio et al. 2018). Notably, accessions ARZM08096 and ARZM08018, which have been recognized as carrying resistance to multiple pathogens (Defacio et al. 2018), exhibit medium-high levels of total phenols, as shown in Table 3a.

Given the increasing extent of soil salinization worldwide (Egea et al. 2023), together with the progression of climate change, the phenotypic variation observed in these landraces may be of considerable value for the development of maize varieties tolerant to salinity and potentially to other abiotic stresses.

## Conclusions

This study provides one of the first comprehensive biochemical characterizations of Argentine maize landraces, integrating morphological and biochemical data with the altitudinal gradient. While previous research has focused on the morphological or genetic diversity of these landraces, the systematic biochemical profiling across multiple compound classes -such as pigments, phenolics, carbohydrates, and antioxidant activity-combined with controlled assessments of stress tolerance, marks a significant advancement in understanding the phenotypic diversity of this germplasm.

Phenotypic variability, including diversity in biochemical composition and stress tolerance, acquires strategic relevance when correlated with genotype, as it contributes to the sustainability and resilience of agricultural systems in the face of climate change and other global challenges (Guzzon et al. 2021; Teixeira et al. 2021). The 19 maize accessions from Northern Argentina evaluated in this study exhibited high variability across the assessed traits at the individual level (Tables 1-3). The combined use of univariate and multivariate analyses revealed that each accession possesses a unique combination of morphological, biochemical, and salinity stress tolerance traits, with overall phenotypic differentiation observed among accessions associated with the altitude of collection. This integrative approach provides new insights into how local environmental conditions contribute to the functional diversification of maize landraces in Argentina.

These results underscore the importance of considering the high variability among accessions and the singularity of each accession when designing both *in situ* and *ex situ* conservation strategies. Such an approach is essential not only for addressing agronomic challenges but also for preserving the cultural heritage associated with maize landraces. Moreover, the phenotypic diversity documented here provides a valuable foundation for future studies, including quantitative trait locus (QTL) mapping and genome-wide association studies (GWAS), aimed at identifying genetic loci linked to agronomically important traits.

## Data Availability Statement

All data supporting the findings of this study are included in the main manuscript, available in the Supplementary materials, or can be obtained directly from the corresponding author upon request.

## Contributions

**D.T.L.:** performed experiments, statistical analyses and data interpretation, wrote the manuscript. **F.D.:** performed statistical analyses and data interpretation, wrote the manuscript. **D.R.A.:** provided the seeds and performed the salt stress tolerance assay. **F.M.:** performed the salt stress tolerance assay. **P.N.B.:** collaborated in providing space and resources for the project. **L.V.V.:** analyzed data, performed data interpretation, provided resources for the project, wrote the manuscript. **D.P.G:** designed experiments, performed statistical analyses and data interpretation, provided resources for the project, wrote the manuscript. All the authors revised the manuscript, provided suggestions and approved the final version of the manuscript.

## Acknowledgments

We would like to express our sincere gratitude to Ana Elisa Font and Macarena Arza for their invaluable assistance in conducting the salinity stress experiments.

