## Supplementary figure 1 for "Southern South American Maize Landraces: A Source of Phenotypic Diversity"

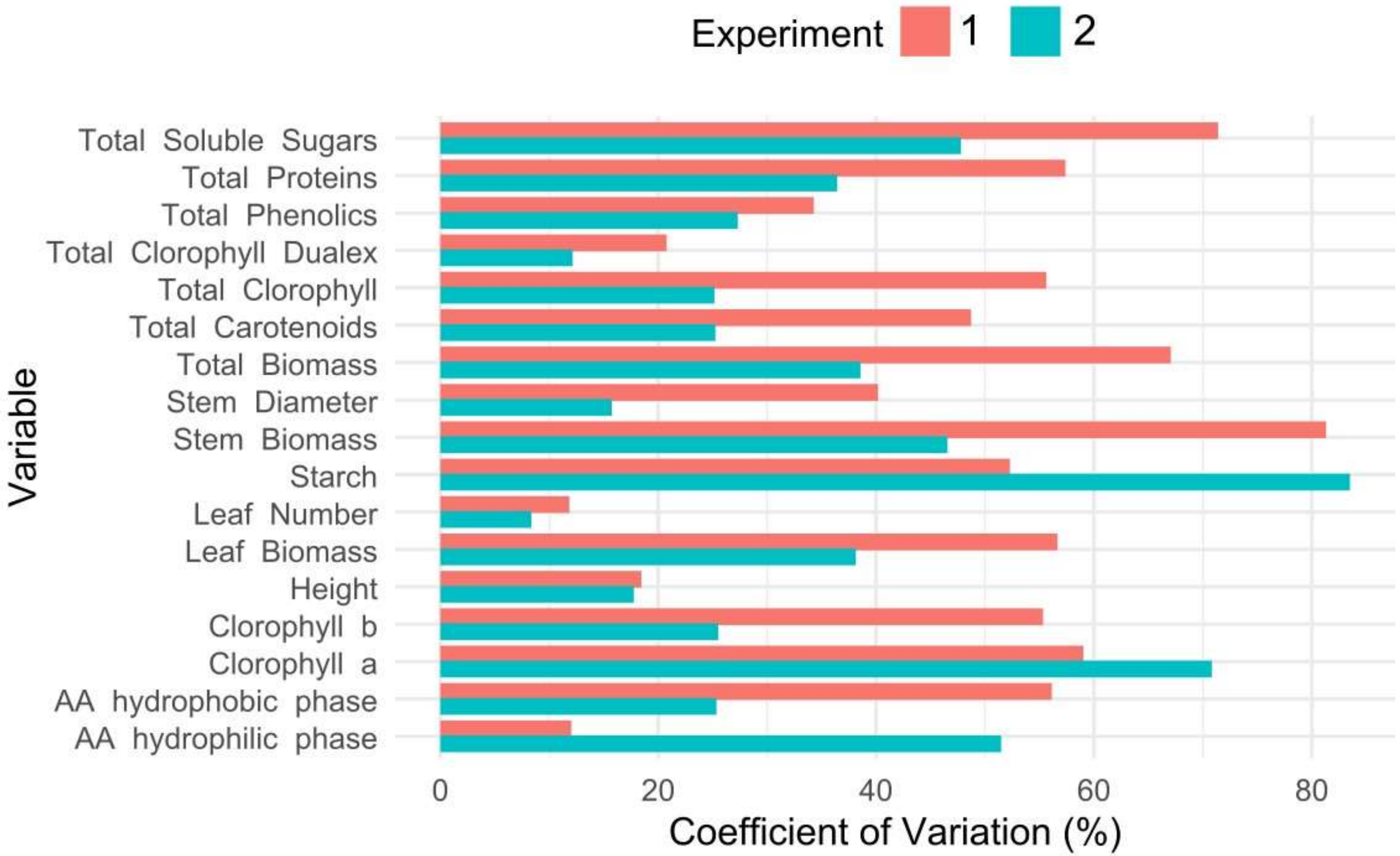


**Supplementary Figure 1.** Coefficients of variation of agromorphological and biochemical traits of Experiment 1 (pink) and 2 (green). AA: Antioxidant Activity.
