## Supplementary figure 2 for "Southern South American Maize Landraces: A Source of Phenotypic Diversity"

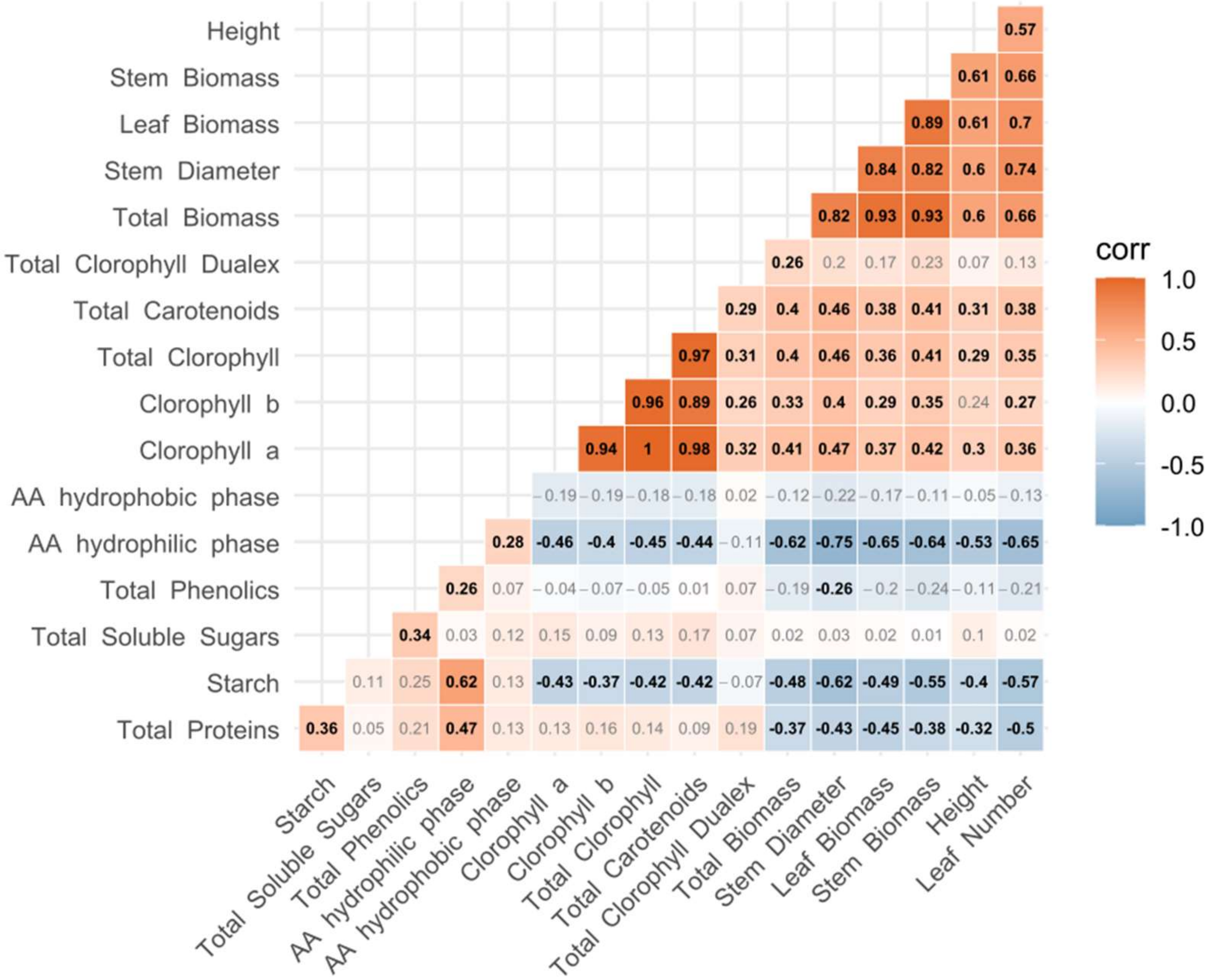


**Supplementary Figure 2. Spearman correlations** between agromorphological and biochemical traits of the 19 accessions, adjusted for multiple testing using the sequential Bonferroni correction. Bold font represents significant results ($p<0.05$).
