## Supplementary figure 3 for "Southern South American Maize Landraces: A Source of Phenotypic Diversity"

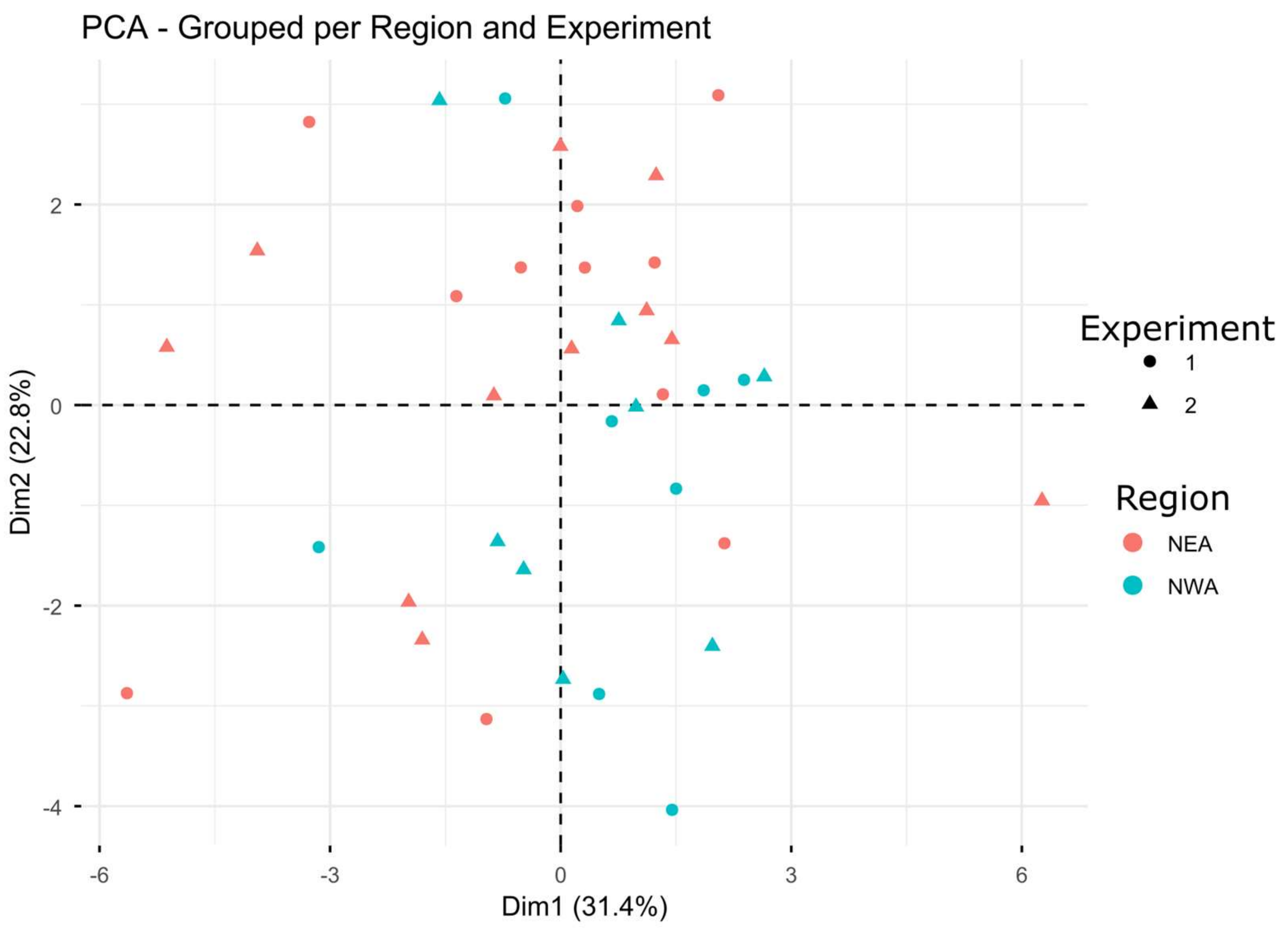


**Supplementary Figure 3. Principal Components Analysis.** Data are classified by Region of origin (NWA: Northwestern Argentina; NEA: Northeastern Argentina) and Experiment (1 and 2). Data were averaged per accession and standardized and centered within each experiment.
