## Supplementary table 1 for "Southern South American Maize Landraces: A Source of Phenotypic Diversity"

**Supplementary table 1.** Characteristics of the accessions evaluated from the "Banco Activo de Germoplasma INTA Pergamino" (BAP; Active Germplasm Bank of the National Institute of Agricultural Technology, Pergamino, Buenos Aires, Argentina). The classification of soils was retrieved from the soil charter of the Instituto Geográfico Nacional (National Geographic Institute, IGN).[a]

| Bank ID | Race | Original collection data | | | | | | |
|---|---|---|---|---|---|---|---|---|
| | | Location | Province | Region | Longitude (º) | Latitude (º) | Altitude (m.a.s.l.) | Soil type |
| **ARZM03042** | Perlita | Ejido Norte, Feliciano | Entre Ríos | NEA | -58.45 | -30.23 | 67 | E |
| **ARZM04013** | Pisingallo | Ista cora, Mercedes | Corrientes | NEA | -58.14 | -29.11 | 53 | M |
| **ARZM04029** | Avatí morotí | Ifrán, Goya | Corrientes | NEA | -58.59 | -29.05 | 66 | M |
| **ARZM04060** | Calchaquí | Corrientes | Corrientes | NEA | -58.50 | -27.3 | 39 | E |
| **ARZM05007** | Pisingallo | Colonia Tarauco, San Ignacio | Misiones | NEA | -55.29 | -27.12 | 235 | A |
| **ARZM05067** | Avatí morotí ti | Campo Las Monjas km 300 | Misiones | NEA | -54.32 | -26.75 | 345 | A |
| **ARZM05069** | Avatí morotí ti | Campo Las Monjas km 300 | Misiones | NEA | -54.32 | -26.75 | 345 | A |
| **ARZM05118** | Avatí morotí | Santo Pipó, San Ignacio | Misiones | NEA | -55.24 | -27.08 | 232 | U |
| **ARZM05120** | Perlita | Hipólito Yrigoyen, San Ignacio | Misiones | NEA | -55.17 | -27.06 | 210 | U |
| **ARZM06060** | Pisingallo | Machagay, 25 de Mayo | Chaco | NEA | -60.03 | -26.56 | 83 | M |
| **ARZM06109** | Avatí morotí | Lapachito, Gral. Donovan | Chaco | NEA | -59.29 | -27.1 | 62 | I |
| **ARZM08018** | Venezolano | El Jardín, Candelaria | Salta | NWA | -65.23 | -26.05 | 1940 | M |
| **ARZM08020** | Pisingallo | La Candelaria, La Candelaria | Salta | NWA | -65.22 | -24.54 | 1000 | I |
| **ARZM08096** | Amarillo de ocho | Cachi, Cachi | Salta | NWA | -66.12 | -25.07 | 2280 | E |
| **ARZM08178** | Dentado blanco | Vallecito | Salta | NWA | -25.22 | -66.20 | 2209 | E |
| **ARZM10036** | Capia blanco | Trancas,Trancas | Tucumán | NWA | -64.17 | -26.13 | 782 | M |
| **ARZM10076** | Venezolano | Los Puestos, Leales | Tucumán | NWA | -65.18 | -27.12 | 322 | M |
| **ARZM10082** | Perla | Amaicha del Valle | Tucumán | NWA | -65.953 | -26.59 | 1700 | E |
| **ARZM12205** | Calchaquí | Colpes, Poman | Catamarca | NWA | -66.13 | -28.06 | 1250 | E |

[a] NEA: Northeastern Argentina. NWA: Northwestern Argentina (NWA). E= Entisols, M=Mollisols, A=Alfisol, U=Ultisols, I=Inceptisols.
