## Supplementary table 2 for "Southern South American Maize Landraces: A Source of Phenotypic Diversity"

**Supplementary table 2.** Agromorphological evaluation of landraces in Experiment 1 and Experiment 2 (Height, Leaf Number, Total Chlorophyll) or combined Experiments 1 and 2 (Biomass and Stem Diameter) classified by Region. Data represent Estimated Marginal Means or EMM (Standard Errors or SE) from Linear Mixed Effects Models, with bold letters indicating significant differences ($p<0.05$), and Coefficients of Variation (CV). N=3-5.[a]

| Variables | Experiment | Region | | | | p-value |
|---|---|---|---|---|---|---|
| | | NEA | | NWA | | |
| | | EMM (SE) | CV (%) | EMM (SE) | CV (%) | |
| **Height (cm)** | 1 | 45.3 (1.73) | 20.37 | 49.7 (1.99) | 14.83 | 0.108 |
| | **2** | **56.8 (2.03)** | **16.91** | **64.1 (2.8)** | **16.53** | **0.048** |
| **Leaf Number** | **1** | **8.07 (0.144)** | **11.5** | **8.79 (0.18)** | **10.63** | **0.002** |
| | 2 | 10.4 (0.142) | 9.1 | 10.5 (0.158) | 7.42 | 0.426 |
| **Total Chlorophyll Dualex® (µg/cm$^2$)** | 1 | 38.9 (1.57) | 22.79 | 39.3 (1.77) | 17.88 | 0.851 |
| | 2 | 39.7 (0.71) | 12.46 | 41.2 (0.77) | 11.57 | 0.159 |
| **Total Biomass (g)** | 1+2 | 57.5 (31.4) | 70.02 | 65.2 (35.9) | 61.59 | 0.886 |
| **Leaf Biomass (g)** | 1+2 | 18.2 (8.77) | 65.02 | 19.7 (9.56) | 58.16 | 0.920 |
| **Stem Biomass (g)** | 1+2 | 23.8 (15.1) | 82.4 | 30.6 (19.5) | 71.56 | 0.804 |
| **Stem Diameter (cm)** | 1+2 | 1.08 (0.621) | 57.94 | 1.2 (0.692) | 52.37 | 0.907 |

[a]NEA: Northeastern Argentina; NWA: Northwestern Argentina.
