## Supplementary table 3 for "Southern South American Maize Landraces: A Source of Phenotypic Diversity"

**Supplementary table 3.** Leaf metabolite content of fully expanded leaves from V12 stage plants in combined Experiments 1 and 2 classified by Region. Data represent Estimated Marginal Means or EMM (Standard Errors or SE) from Linear Mixed Effects Models, with bold letters indicating significant differences (p<0.05), and Coefficients of Variation (CV). N=3-5.[a]

| Variables | Region | | | | p-value |
|---|---|---|---|---|---|
| | NEA | | NWA | | |
| | MM (SE) | CV (%) | MM (SE) | CV (%) | |
| **Total Proteins (mg/g)** | 1.45 (0.298) | 70.86 | 1.52 (0.304) | 60.67 | 0.884 |
| **Total Soluble Sugars (mg/g)** | 11.2 (0.638) | 68.82 | 13.6 (0.912) | 54.11 | 0.166 |
| **Starch (mg/g)** | 11.79 (13.58) | 83.64 | 8.53 (9.84) | 107.86 | 0.861 |
| **Total Chlorophyll (g/g)** | 1.943 (0.892) | 51.19 | 0.987 (0.455) | 82.11 | 0.404 |
| **Chlorophyll a (g/g)** | 1.55 (0.622) | 51.48 | 0.845 (0.341) | 76.04 | 0.394 |
| **Chlorophyll b (g/g)** | 0.495 (0.309) | 88.58 | 0.914 (0.571) | 74.11 | 0.558 |
| **Total Carotenoids (g/g)** | 0.46 (0.347) | 114.44 | 0.957 (0.724) | 82.53 | 0.563 |
| **Total Phenolics (mg GA/g)** | 2.01 (0.142) | 25.7 | 2.15 (0.156) | 37.89 | 0.567 |
| **Antioxidant Activity (Hydrophilic phase) (mg GA/g)** | 0.0717 (0.0539) | 59.46 | 0.0673 (0.0506) | 65.7 | 0.958 |
| **Antioxidant Activity (Hydrophobic phase) (mg GA/g)** | 0.0732 (0.00737) | 60.48 | 0.0749 (0.00861) | 71.96 | 0.892 |

[a]NEA: Northeastern Argentina; NWA: Northwestern Argentina. GA= Gallic Acid.
