## Supplementary table 4 for "Southern South American Maize Landraces: A Source of Phenotypic Diversity"

**Supplementary table 4.** Scores for phenotypic data obtained in the Redundancy Analysis (RDA). Bold letters indicate the major contributors to the RDA axes (scores >0.8 or < -0.8).[a]

| Phenotypic variables | RDA1 | RDA2 |
|---|---|---|
| **Total Proteins** | -0.6058316 | **-0.82100179** |
| **Total Soluble Sugars** | **1.3741597** | -0.09691631 |
| **Starch** | **-1.2153123** | **-1.02543119** |
| **Total Chlorophyll** | **-1.3340648** | 0.08019263 |
| **Chlorophyll a** | **-1.8927774** | 0.75048773 |
| **Chlorophyll b** | **0.8019153** | -0.47289632 |
| **Total Carotenoids** | **0.9404102** | -0.14568723 |
| **Total Phenolics** | **1.4373382** | 0.03379227 |
| **AA Hydrophilic Phase** | **-1.5260978** | -0.07209119 |
| **AA Hydrophobic phase** | 0.1019801 | 0.77521652 |
| **Total Chlorophyll Dualex®** | **0.8608896** | **-2.25679819** |
| **Total Biomass** | 0.2677883 | **1.09507074** |
| **Stem Biomass** | 0.517789 | **0.80130035** |
| **Leaf Biomass** | 0.2213693 | **1.08258775** |
| **Stem Diameter** | 0.523741 | **2.27343712** |

[a]AA: Antioxidant Activity.
